# Meiotic resetting of the cellular Sod1 pool is driven by protein aggregation, degradation, and transient LUTI-mediated repression

**DOI:** 10.1101/2022.06.28.498006

**Authors:** Helen M. Vander Wende, Mounika Gopi, Megan Onyundo, Claudia Medrano, Temiloluwa Adanlawo, Gloria A. Brar

## Abstract

Gametogenesis requires packaging of the cellular components needed for the next generation. In budding yeast, this process includes degradation of many mitotically stable proteins, followed by their resynthesis. Here, we show that one such case—Superoxide dismutase 1 (Sod1), a protein that commonly aggregates in human ALS patients—is regulated by an integrated set of events, beginning with the formation of pre-meiotic Sod1 aggregates. This is followed by degradation of a subset of the prior Sod1 pool and clearance of Sod1 aggregates. As degradation progresses, Sod1 protein production is transiently blocked during mid-meiotic stages by transcription of an extended and poorly translated *SOD1* mRNA isoform, *SOD1^LUTI^*. Expression of *SOD1^LUTI^* is induced by the Unfolded Protein Response, and it acts to repress canonical *SOD1* mRNA expression. *SOD1^LUTI^* is no longer expressed following the meiotic divisions, enabling a resurgence of canonical mRNA and synthesis of new Sod1 protein such that gametes inherit a full complement of this important enzyme that is essential for gamete viability. Altogether, this work reveals meiosis to be an unusual cellular context in which Sod1 levels are tightly regulated. Our findings also suggest that further investigation of Sod1 during yeast gametogenesis could shed light on conserved aspects of its aggregation and degradation that could have implications for our understanding of human disease.

## INTRODUCTION

The transformation of precursor cells into gametes by meiosis and gametogenesis is responsible for determining which cellular material is passed on to the next generation. In the budding yeast *Saccharomyces cerevisiae*, this complex differentiation program is driven by tightly regulated changes in protein synthesis for almost every annotated gene, including many with no established roles in meiosis or gamete formation (Brar et al., 2012; Cheng & Otto et al., 2018). For many genes, this regulation is achieved via transcript toggling between expression of a canonical mRNA isoform and a poorly translated <u>L</u>ong <u>U</u>ndecoded <u>T</u>ranscript <u>I</u>soform (LUTI). LUTIs are 5′-extended transcripts containing competitive upstream ORFs (uORFs) whose translation repress translation of the main ORF (Chen et al., 2017; Tresenrider et al., 2021). Transcription of LUTIs interferes in cis with the downstream transcription start site that drives the canonical, translatable mRNA isoform. This noncanonical regulation is common during meiosis in budding yeast, regulating approximately 8% of genes, and is a core part of the ER Unfolded Protein Response (UPR^ER^) that long went unrecognized (Cheng and Otto et al., 2018; Van Dalfsen et al., 2018). LUTI-like regulation has also been found to control gene expression in diverse organisms, including human cells, flies, and plants (Corbin and Maniatis, 1989; Moseley et al., 2002; Sehgal et al., 2008; Hollerer et al., 2019; Jorgensen et al., 2020).

The functional significance of LUTI-based regulation has been shown for the kinetochore gene, *NDC80* (Chen et al., 2017; Chia et al., 2017), for which dynamic modulation of protein levels has a known role in chromosome segregation during meiosis (Miller and Ünal et al., 2012). However, we previously identified hundreds of genes that seem to be regulated in this manner, including so-called housekeeping genes which are thought to be constitutively expressed (Cheng & Otto et al., 2018). An example is *SOD1*, which encodes the highly abundant antioxidant enzyme Superoxide dismutase 1, a Cu-Zn superoxide dismutase that converts superoxide radicals into hydrogen peroxide and molecular oxygen (McCord and Fridovich, 1969). Sod1 is highly conserved from yeast to humans, and the human *SOD1* gene can complement loss of the yeast gene (Corson et al., 1998). Human SOD1 (hSOD1) has been studied extensively due to its involvement in familial cases of Amyotrophic Lateral Sclerosis (fALS). Over one hundred unique point mutations in the *SOD1* gene have been identified in ALS patients (Saccon et al., 2013), many of which increase the propensity of hSOD1 to form aggregates, particularly within tissues of the nervous system (Watanabe et al., 2001). Despite the discovery of Sod1’s association with ALS nearly 30 years ago (Rosen et al., 1993), its precise role in disease progression remains unclear. Some models propose that Sod1 aggregates are toxic, whereas others propose that they may be protective (Gill et al., 2019). The links between Sod1’s aggregation, toxicity, and degradation have been difficult to mechanistically assess, in part due to the complex nature of the contexts in which aggregation has been observed (Pansarasa et al., 2018; Di Gregorio and Duennwald, 2018).

Sod1 is inherently highly stable in its fully folded and metalated state, even remaining enzymatically active at 90^°^C (Hallewell et al., 1991) and—most remarkably— in tissue from a 3,000-year-old mummy (Weser et al., 1989). During exponential mitotic growth, yeast Sod1 is thought to be decreased not by active degradation but rather by passive dilution, along with 85% of the proteome (Christiano et al., 2014). We investigated the impact of LUTI regulation on Sod1 protein levels during gametogenesis and found that it drives transient inhibition of new Sod1 synthesis during the meiotic divisions, followed by rapid Sod1 protein repopulation. This coincides with degradation of preexisting Sod1, which begins prior to meiosis and follows the pervasive natural aggregation of wild-type protein. These findings reveal a complex and coordinated gene regulatory program during gametogenesis that achieves depletion of preexisting protein and replenishment of the Sod1 protein pool that is passed on to the next generation. Moreover, this work reveals yeast meiosis as a useful system for studying the differential regulation of both wild-type and ALS mutant Sod1 protein.

## RESULTS

### An alternative transcript isoform is expressed from the *SOD1* locus during meiosis

A previous global study from our lab identified a high degree of regulation for Sod1 during meiosis in budding yeast, with its translation and protein levels dropping in mid-meiosis and rising again as gametes (spores) are formed (Cheng and Otto et al., 2018). We also found *SOD1* to be one of nearly 400 genes that showed signatures of LUTI-based regulation (Cheng and Otto et al., 2018), including an unexpectedly poor correlation between mRNA and protein levels over a 12-hour meiotic time course (Figure 1A). Analysis of the *SOD1* locus in these global datasets revealed that during mitotic (vegetative) growth and early in meiosis, budding yeast cells expressed the expected canonical *SOD1* transcript (*SOD1^canon^*.), which is roughly 600 nucleotides long (Figure 1B). During the meiotic divisions, however, mRNA-seq read density extended from this canonical locus to a region 1.6 kb upstream of the canonical transcription start site (TSS), which appeared to be a result of production of an extremely 5′ extended mRNA isoform (Figure 1B). The presence of this elongated transcript was correlated with translation of an upstream open reading frame (uORF) that is housed in the extended 5′ transcript region, as well a decrease in ribosome footprint density mapping to the *SOD1* ORF (Figure 1B). These signatures of transcription and translation suggested that a LUTI was expressed from the *SOD1* locus during the meiotic divisions (Cheng and Otto et al., 2018). We became interested in studying this case further because Sod1 protein is thought to be constitutively and abundantly expressed, making temporary repression of its synthesis during meiosis surprising. Furthermore, whereas mitotically dividing cells lacking Sod1 show reduced fitness (Figure S1A), Sod1 is completely essential for the production of viable gametes (Figure 1C).

**Figure 1.**
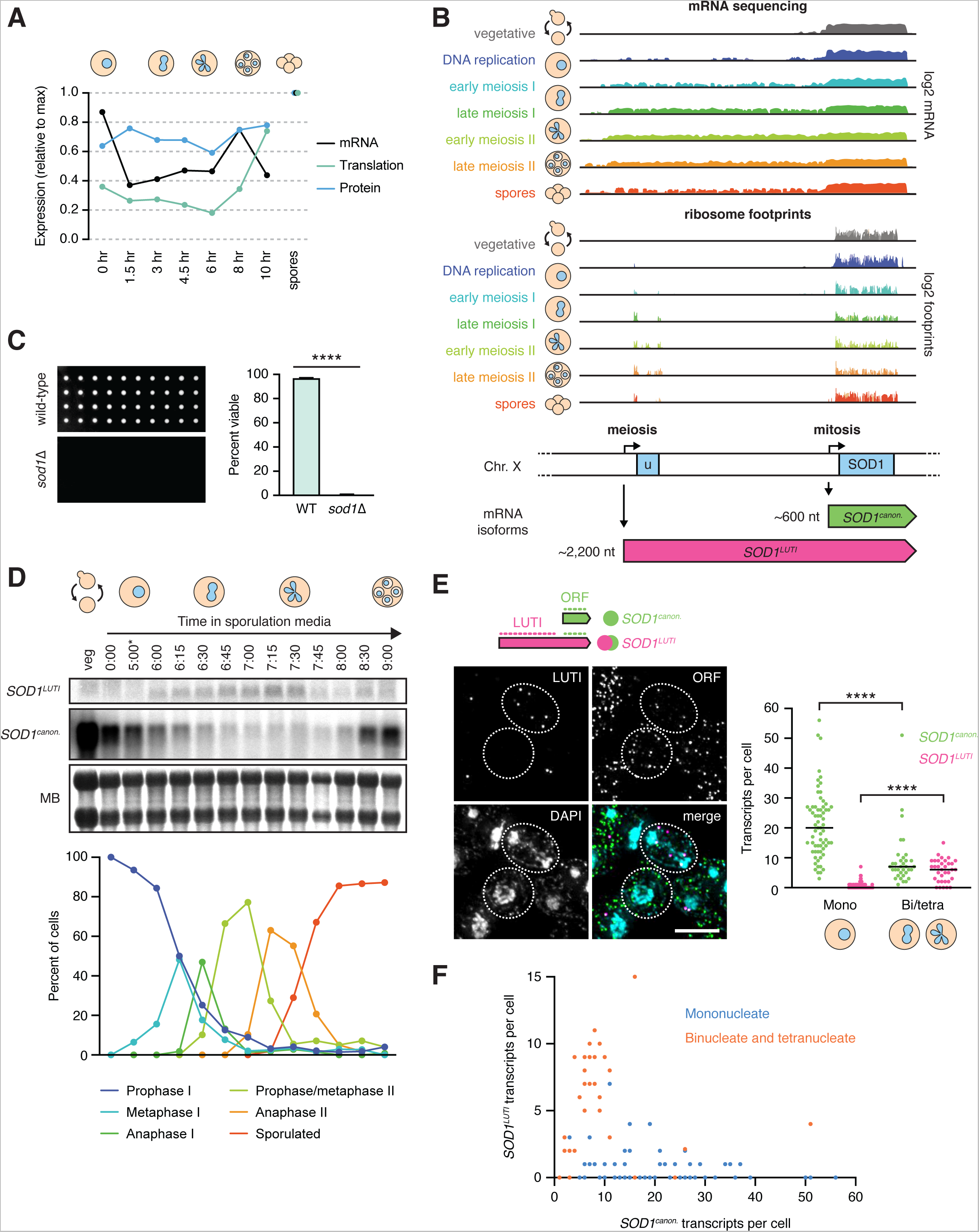
An alternative transcript isoform is expressed from the *SOD1* locus during meiosis. (A) Matched relative expression of *SOD1* mRNA, translation, and protein (Cheng and Otto et al., 2018). All values are normalized to max expression (spores). (B) mRNA-sequencing (top) and ribosome profiling (bottom) reads mapping to the *SOD1* locus of the *Saccharomyces cerevisiae* genome (on Chr. X) in vegetative growth and throughout a meiotic time course (Brar et al., 2012). During the meiotic divisions, ribosome footprints show translation of a uORF of 293 nucleotides. (C) Viability of spores derived from wild-type control and homozygous *sod1*Δ cells dissected on rich media (YEP + 2% dextrose) and incubated at 30°C for 48 hours. Each column represents four spores from the same tetrad, and quantification represents the average viability of 20 tetrads of each genotype (biological triplicate). (D) Northern blot probing for *SOD1* mRNA (top) throughout a Ndt80-synchronized meiotic time course (MB = methylene blue, \**pGAL*-*NDT80* release at 5 hours) and matched tubulin immunofluorescence (bottom; at least 100 cells counted per time point). (E) Single molecule RNA fluorescence *in situ* hybridization (smRNA-FISH) probing for *SOD1* mRNAs using two sets of probes (scale bar = 5 µm). Quantification of smRNA-FISH data in mononucleate (Mono) vs. binucleate/tetranucleate (Bi/tetra) cells shows a significant drop in *SOD1^canon^*. levels (Mann-Whitney U = 340, P <0.0001) and a significant increase in *SOD1^LUTI^* levels (Mann-Whitney U = 275.5, P < 0.0001) (mononucleate n=66, binucleate/tetranucleate n=34). (F) smRNA-FISH quantification of *SOD1^canon^*. (X axis) vs. *SOD1^LUTI^* (Y axis) transcripts per cell. For quantification in E and F, *SOD1^LUTI^* transcripts were defined as colocalized foci of ‘LUTI’ and ‘ORF’ probe sets (‘LUTI’-only foci, representing ∼30% of LUTI probe signal, were excluded from analysis).

Our mRNA-seq data suggested that a 5′ extended *SOD1* transcript (*SOD1^LUTI^*) of approximately 2.2 kilobases (kb) in length was transiently produced during meiosis, but these data could not preclude the possibility that the mRNA read density represented an adjacent transcript. To test whether *SOD1^LUTI^* is expressed as a continuous mRNA, we performed northern blotting using a probe that hybridized to a sequence within the *SOD1* ORF to detect both canonical (*SOD1^canon^*.) and LUTI (*SOD1^LUTI^)* transcripts. To increase temporal resolution of *SOD1* mRNA expression changes, we used a strain expressing both *pGAL-NDT80* and a *GAL4-ER* trans-activator, allowing us to arrest cells in prophase I until the addition of β-estradiol, which results in highly synchronous progression through subsequent meiotic stages (Carlile and Amon, 2008). During the meiotic divisions, approximately from metaphase I to anaphase II as assessed by tubulin immunofluorescence, *SOD1^canon.^* expression decreased dramatically, and this decrease corresponded in timing with appearance of a higher band that represents *SOD1^LUTI^* (Figure 1D). After both meiotic divisions occur, LUTI expression decreased, and canonical mRNA was restored. This “transcript toggling” is a hallmark of LUTI-based regulation and suggests that *SOD1^LUTI^* blocks *SOD1^canon^*. expression through transcriptional interference (Chen et al., 2017; Chia et al., 2017; Cheng and Otto et al., 2018; Chia et al., 2021; Tresenrider et al., 2021).

We further validated the expression of *SOD1^LUTI^* by single molecule RNA fluorescence *in situ* hybridization (smRNA-FISH; Raj et al., 2008; Chen et al., 2018). Using two fluorescently labeled probe sets that hybridize to either shared sequences within the *SOD1* ORF or LUTI-specific sequences, we visualized individual *SOD1^canon^*. and *SOD1^LUTI^* transcripts in single cells. To examine mRNA expression during meiosis, we fixed cells after 6 hours in sporulation media (SPO) and counted transcript levels in mononucleate and binucleate/tetranucleate cells (Figure 1E). Signal representing LUTI-specific regions colocalized with *SOD1* ORF regions in ∼70% of cases (173/241 foci), and these were the foci that we scored as representing LUTIs. Consistent with our population-based measurements (Figure 1B, 1D), in mononucleate cells which have yet to begin the process of chromosome segregation, *SOD1^canon^*. was expressed almost exclusively. In contrast, binucleate and tetranucleate cells (representing cells late in the first meiotic division and after) showed a significant reduction in *SOD1^canon^*. expression, which coincided with an increase in *SOD1^LUTI^* expression (Figure 1E). On a single-cell level, we observed an inverse relationship between LUTI and canonical transcript abundance, a defining feature of LUTI-based regulation (Figure 1F; Chen et al., 2017; Chia et al., 2017; Tresenrider et al., 2021; Cheng and Otto et al., 2018; Van Dalfsen et al., 2018; Hollerer et al., 2019). Taken together, our northern blot and smRNA-FISH data support the existence of *SOD1^LUTI^*, a meiotic mRNA isoform that is associated with reduced canonical *SOD1* mRNA expression.

### *SOD1^LUTI^* expression depends on the meiotic program and is sufficient to downregulate canonical *SOD1* mRNA

Because *SOD1^LUTI^* transcription is coincident with the meiotic nuclear divisions, we suspected that its expression was downstream of Ndt80, a transcription factor responsible for the induction of a large set of mid-meiotic genes and thus meiotic progression past prophase (Xu et al., 1995; Chu et al., 1998). After a five-hour incubation in sporulation media to synchronize *pGAL*-*NDT80* cells in prophase, cultures were split and treated with β-estradiol (“*NDT80* release”) or ethanol (vehicle control; “*NDT80* block”). RT-qPCR analysis revealed that *SOD1^LUTI^* expression peaked between 2 to 3 hours after β-estradiol treatment (7-8 total hours in SPO) and was not observed in control samples (Figure 2A, top). Comparison of *SOD1^canon^*. by northern blotting in the presence or absence of *NDT80* expression showed that *SOD1^canon^*. levels decreased in both cases (Figure 2A, bottom), but were lower when *NDT80* was expressed, coincident with expression of *SOD1^LUTI^* (Figure 2A, top). These data demonstrate that *SOD1^LUTI^* acts downstream of *NDT80* to repress *SOD1^canon^*. during meiosis. Based on the delayed timing of *SOD1^LUTI^* relative to known direct Ndt80 targets (Cheng and Otto et al., 2018) and the lack of characterized Ndt80 binding sites upstream of the LUTI TSS (Chu et al., 1998), it is likely that *SOD1^LUTI^* is not induced directly by Ndt80. We noted that the robust resurgence of canonical mRNA seen after the meiotic divisions is absent in cells lacking Ndt80 expression (Figure 2A), indicating that both the strong decrease and eventual reappearance of translatable *SOD1* mRNA depends on the meiotic program.

**Figure 2.**
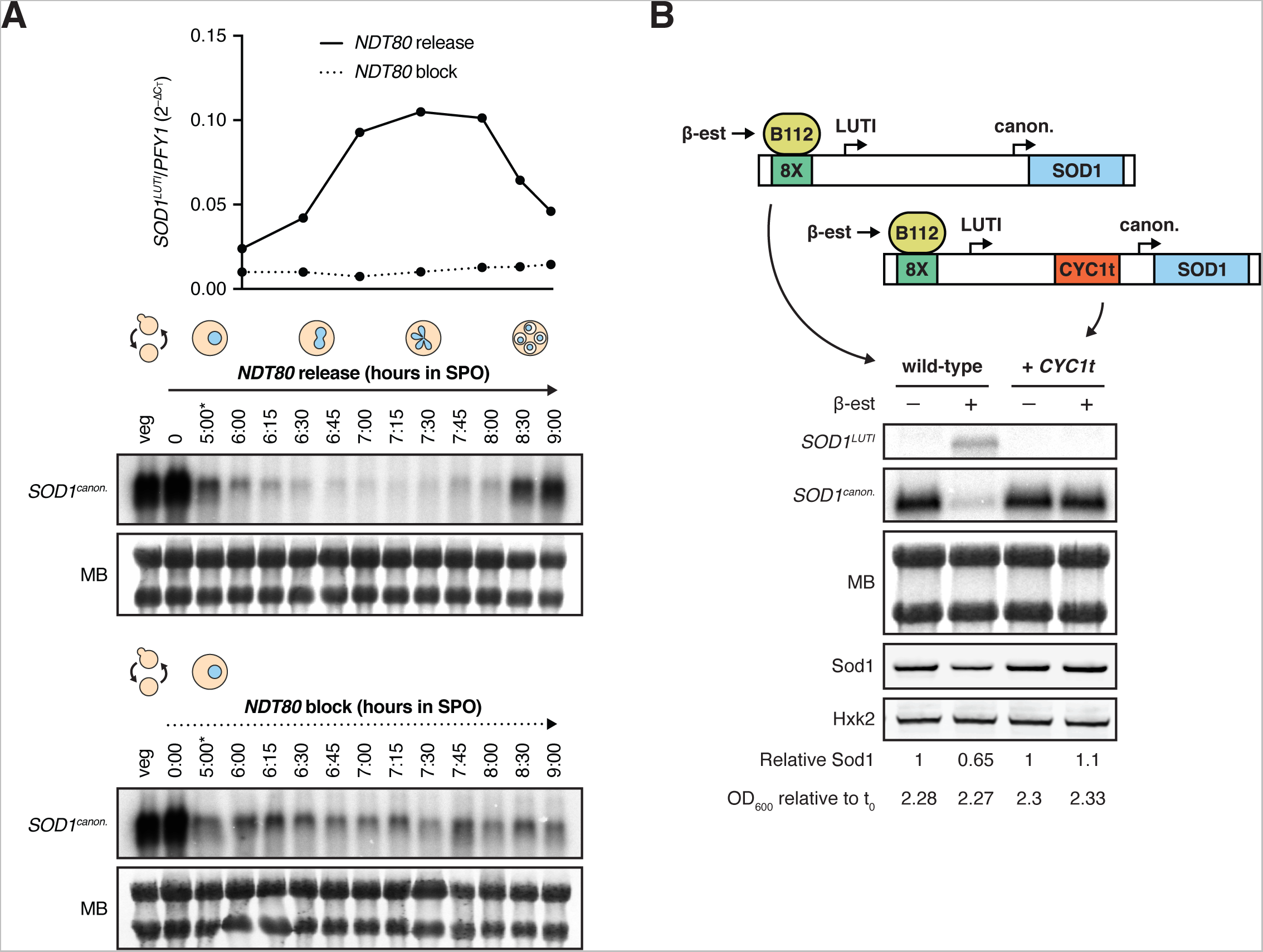
*SOD1^LUTI^* expression depends on the meiotic program and is sufficient to downregulate canonical *SOD1* mRNA. (A) RT-qPCR analysis of *SOD1^LUTI^* (top) and northern blots probing for *SOD1^canon^*. (bottom) in the presence or absence of *NDT80* (MB = methylene blue; \**pGAL*-*NDT80* release at 5 hours). (B) Northern blot and SDS-PAGE immunoblot probing for *SOD1* mRNA and Sod1 protein upon mitotic overexpression of *SOD1^LUTI^* via an inducible lexA/lexO system. To disrupt the LUTI, a transcriptional terminator (*CYC1t*) was inserted between the TSSs. Samples shown were harvested two hours post-treatment with either vehicle control (100% ethanol) or 30 nM β-estradiol. Immunoblot quantification was performed by normalizing to Hexokinase (Hxk2) expression and ‘Relative Sod1’ refers to expression of Sod1 in the treated vs. untreated sample for each genotype.

To test whether mitotic overexpression of *SOD1^LUTI^* is sufficient to reduce *SOD1^canon.^,* we integrated an 8lexO array just upstream of the LUTI TSS, as determined by mRNA-seq and transcript leader (TL-seq) data (Brar et al., 2012; Chia et al., 2021), in a strain containing an inducible lexA trans-activator (B112) to allow conditional overexpression of *SOD1^LUTI^* (Ottoz et al., 2014). Indeed, during vegetative exponential growth, overexpression of *SOD1^LUTI^* results in a robust decrease in *SOD1^canon.^* (Figure 2B). Previous work has found that the characterized transcriptional repression associated with LUTI-based interference relies on transcription from the upstream LUTI TSS through the canonical promoter (Chia et al., 2017). To test whether this was the case for *SOD1*, we used CRISPR-Cas9 (Jinek et al., 2012; Anand et al., 2017) to insert a transcriptional terminator sequence from the *CYC1* gene (*CYC1t*) prior to the canonical promoter and, consistently, observed no decrease in *SOD1^canon.^* in this case (Figure 2B). Given these results, we conclude that production of *SOD1^LUTI^* is both necessary and sufficient to drive down canonical *SOD1* transcript levels.

### The UPR^ER^ drives *SOD1^LUTI^* expression

The UPR^ER^ is naturally and transiently activated during meiosis (Brar et al., 2012; Cheng and Otto et al., 2018), reflected by translation of the conserved UPR^ER^ transcription factor Hac1, the ortholog of metazoan XBP1. We noted that *SOD1^LUTI^* expression begins approximately when UPR^ER^ activation occurs and subsides when UPR^ER^ activation is no longer seen (Figure 3A). Furthermore, two putative Hac1 binding sites, or unfolded response elements (UPREs; Mori et al., 1992; Kohno et al., 1993; Fordyce et al., 2012), exist upstream of the *SOD1^LUTI^* TSS (Figure 3A). We therefore hypothesized that LUTI expression may be induced by UPR^ER^ activation. To test this, we treated mitotic cells during vegetative exponential growth with dithiothreitol (DTT) or tunicamycin (Tm), two drugs commonly used to activate of the UPR^ER^. We found that treatment with either drug drove *SOD1^LUTI^* expression as assessed by smRNA-FISH in wild-type cells (Figures 3B, 3C). This effect was largely dependent on the presence of Hac1 (Figures 3B, 3C), indicating that the conserved Hac1/XBP1 branch of the UPR^ER^ drives expression of *SOD1^LUTI^.* We also found that this regulation was not general to cellular stress, as no *SOD1^LUTI^* production resulted from other forms of oxidative stress or heat shock (Figure S2A).

**Figure 3.**
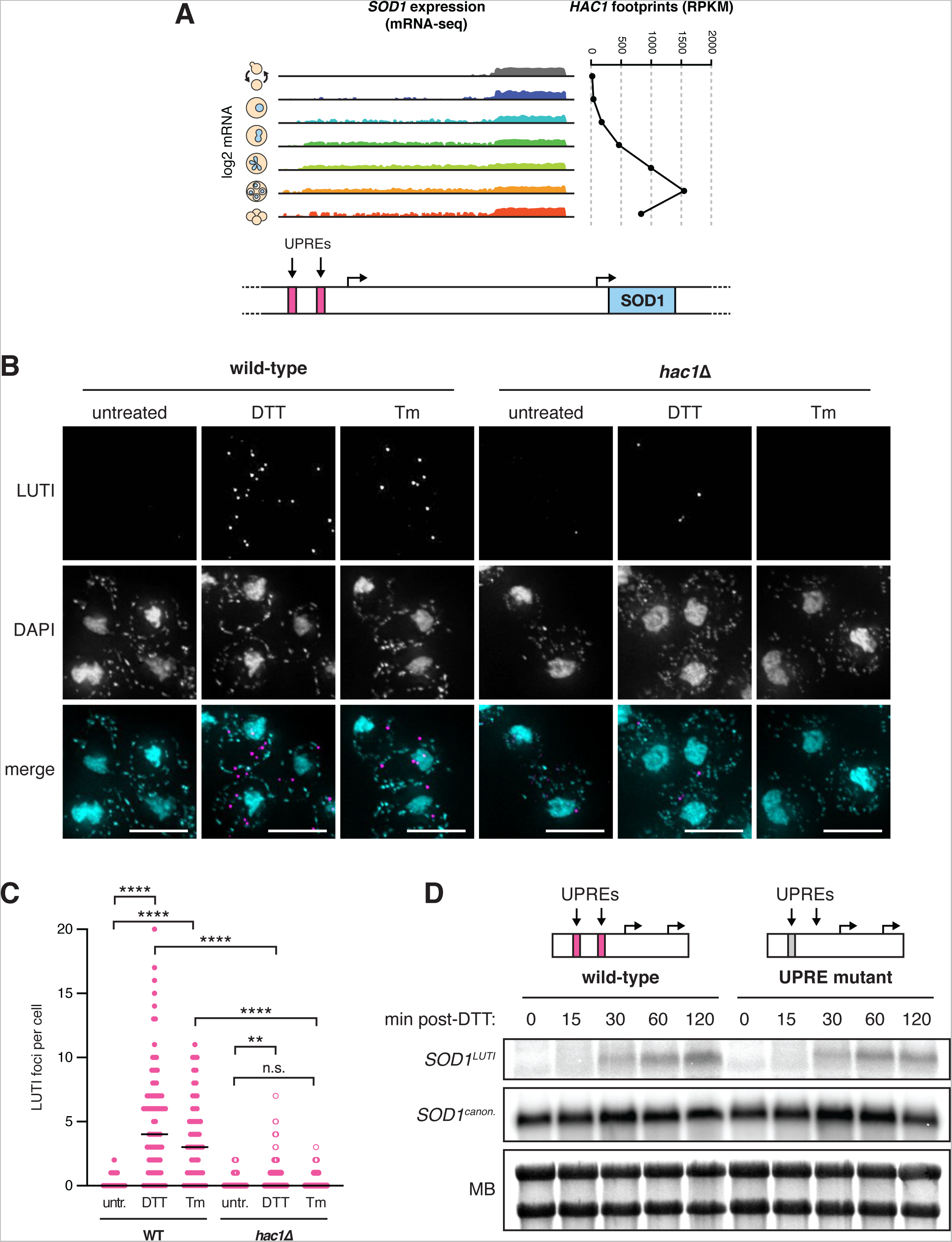
The UPR^ER^ drives *SOD1^LUTI^* expression. (A) mRNA-seq reads mapping to the *SOD1* locus during meiosis (Brar et al., 2012) next to matched *HAC1* translation data (ribosome footprints). (B) smRNA-FISH using the *SOD1^LUTI^*-specific probe set in wild-type and *hac1*Δ vegetative cells fixed 1 hour after treatment with 5 mM DTT or 2 µg/mL Tm (scale bars = 5 µm). (C) Quantification of smRNA-FISH LUTI foci from experiment shown in 3B. For wild-type cells, significant increases in LUTI foci were observed with both DTT (Mann-Whitney U = 695.5, P < 0.0001) and Tm (Mann-Whitney U = 599.5, P < 0.0001). For *hac1*Δ cells, DTT still resulted in a significant increase in LUTI foci (Mann-Whitney U = 1493, P = 0.0038), but Tm did not (Mann-Whitney U = 972, P = 0.5042). The differences between wild-type and *hac1*Δ cells treated with DTT (Mann-Whitney U = 1117, P < 0.0001) and Tm (Mann-Whitney U = 354.4, P < 0.0001) were also significant. Cell counts: WT untr. n = 74, WT DTT n = 86, WT Tm n = 56, *hac1*Δ untr. n = 52, *hac1*Δ DTT n = 77, *hac1*Δ Tm n = 40. (D) Northern blot probing for *SOD1* mRNAs in wild-type and UPRE mutant vegetative exponential cultures 0-120 minutes after 5 mM DTT treatment.

To determine the potential importance of the UPREs located upstream of the LUTI TSS, we used Cas9 to delete the proximal UPRE. Due to its overlap with a gene on the opposite strand (*ECM27*), we could not delete the distal UPRE and instead used Cas9 to scramble the sequence to abolish the predicted UPRE while maintaining the coding sequence of *ECM27*. When cells were treated with DTT during vegetative exponential growth, LUTI expression was still observed by northern blotting (Figure 3D) and RT-qPCR (Figure S2B) in the absence of both UPREs. From these data, we concluded that these predicted UPREs are not essential for *SOD1^LUTI^* expression. This result could indicate that either *SOD1^LUTI^* production is indirectly dependent on Hac1, or that it is dependent on Hac1 binding to elements other than these predicted UPREs. There is precedent for Hac1-dependent transcriptional activation through DNA motifs that have yet to be defined, as approximately half of known Hac1-dependent UPR targets identified by drug-based activation do not contain predicted UPREs within 1 kb of their TSS (Travers and Patil et al., 2000; Patil et al., 2004; Van Dalfsen et al., 2018).

### *SOD1^LUTI^* expression and abatement modulate Sod1 levels

What impact does this transcript toggling have on meiotic Sod1 protein levels? The timing of *SOD1^LUTI^* expression during meiosis (Figure 1D), which drives loss of the canonical and translatable transcript (Figure 2B), was correlated with a steady decrease in Sod1 protein to less than half of vegetative levels after 8 hours in sporulation media (Figure 4A). Furthermore, forced expression of *SOD1^LUTI^* in mitotic cells reduces Sod1 protein levels to ∼65% of wild-type levels, consistent with severely reduced new protein synthesis and dilution of the pre-existing pool by cellular division (Figure 2B). Following the meiotic divisions, Sod1 levels increased with timing that mirrored the return of *SOD1^canon.^* transcript levels (Figures 1B, 1D), with Sod1 protein returning to its early meiotic levels within 4 hours and approaching vegetative abundance levels by 24 hours, when spores are fully formed (Figure 4A). The timed loss and resurgence of Sod1 protein during meiosis that we observed using endogenously tagged Sod1-3V5 mirrored mass spectrometry results in a strain with untagged Sod1 protein (Figure 1A; Cheng and Otto et al. 2018), arguing that the 3V5 tag did not interfere with the normal regulation of Sod1 protein abundance in meiosis. Additionally, we observed a similar pattern of total Sod1 during meiosis in 8M urea-denatured lysates (Figure S3A), arguing that SDS-solubility changes in Sod1 did not drive these trends.

**Figure 4.**
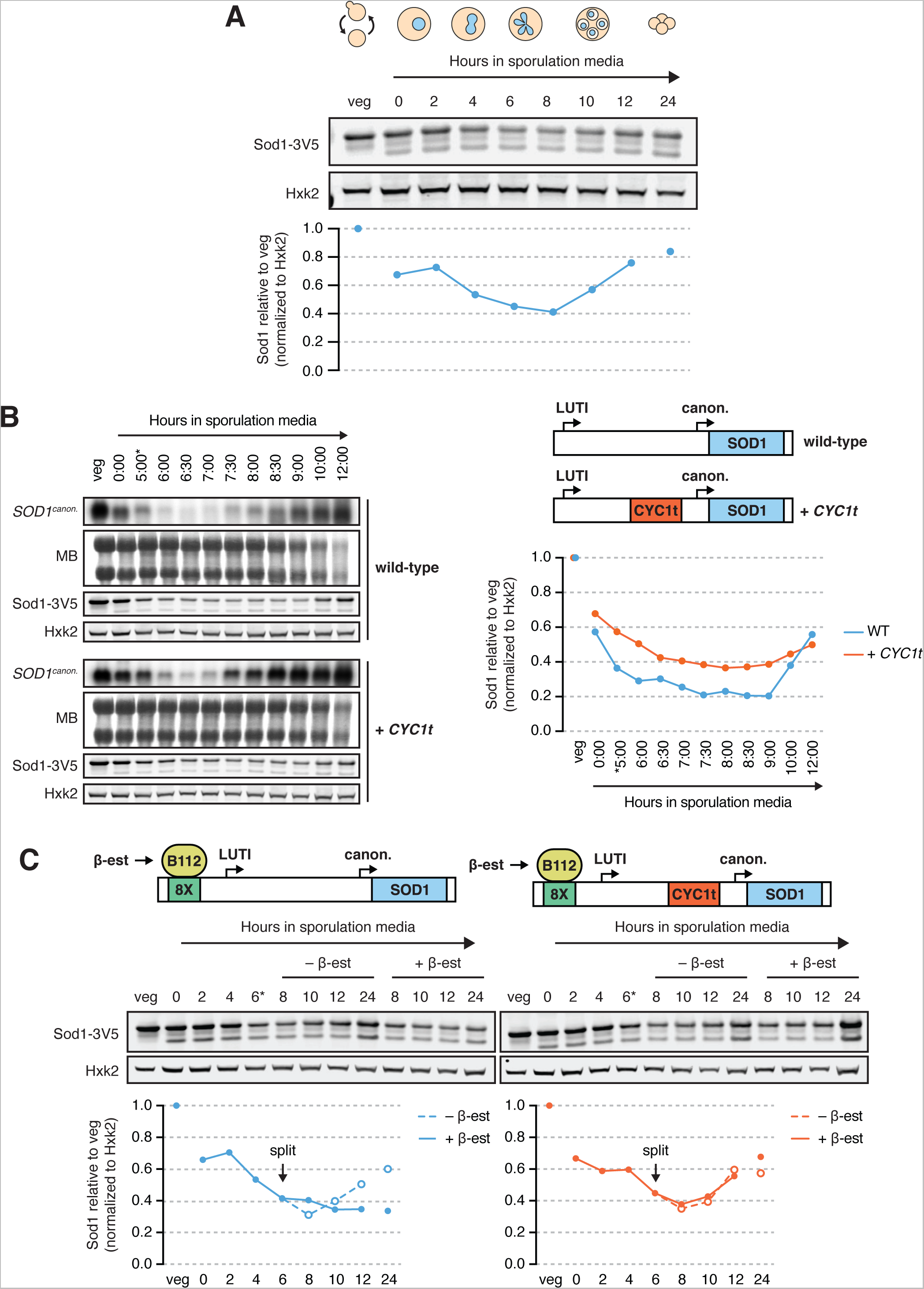
*SOD1^LUTI^* expression and abatement modulate Sod1 levels. (A) SDS-PAGE and immunoblotting for Sod1 throughout a meiotic time course (quantification shown below). (B) Northern blots and SDS-PAGE immunoblots surveying *SOD1* mRNA and Sod1 protein levels in Ndt80-synchronized cells with or without LUTI expression (MB = methylene blue, \**pGAL*-*NDT80* release at 5 hours). To disrupt the LUTI, a transcriptional terminator (*CYC1t*) was inserted between the TSSs. Immunoblot quantification (right) represents one replicate of the data in Figure S3B. (C) Sod1 protein levels in the presence or absence of LUTI overexpression mid-meiosis. To overexpress the LUTI, meiotic cultures of lexA/lexO strains (also used in Figure 2B) were split after 6 hours in sporulation media and treated with either ethanol (vehicle control) or 30 nM β-estradiol (quantification shown below).

To directly test causality of *SOD1^LUTI^* expression on the loss of Sod1 protein in mid-meiosis, we inserted the *CYC1* transcription termination sequence (*CYC1t*) between the distal and proximal TSSs at the endogenous *SOD1* locus to prematurely terminate *SOD1^LUTI^* transcripts prior to the canonical TSS. Northern blotting of LUTI-disrupted cells demonstrated increased *SOD1^canon.^* abundance relative to wild-type during the period in which *SOD1^LUTI^* is typically expressed (Figure 4B). Some decrease in canonical transcript is seen even when the LUTI is disrupted by *CYC1t* insertion and the cause of this decrease is unknown, but a stronger and more sustained decrease in canonical *SOD1* mRNA is seen when full-length *SOD1^LUTI^* is transcribed, demonstrating that *SOD1^LUTI^* transcription through the *SOD1^canon.^* TSS drives the bulk of the transient decrease in canonical mRNA that occurs during meiosis. Examination of Sod1 protein levels in these strains revealed LUTI-disrupted (+ *CYC1t*) cells to show a significant increase in Sod1 protein during mid-meiosis compared to wild-type controls (Figure 4B, Figure S3B), consistent with the hypothesis that the increase in *SOD1^canon^*. in the absence of LUTI-based repression leads to translation of new Sod1 protein.

After the meiotic divisions, *SOD1^LUTI^* expression ceases and *SOD1^canon^*. levels increase (Figures 1B, 1D, 4B), allowing for synthesis of new Sod1 protein concomitant with spore formation (Figures 1A, 4A, 4B). To test whether loss of LUTI production is needed for this resurgence of Sod1 protein levels, 8XlexO-driven *SOD1^LUTI^* was induced at 6 hours after transfer to sporulation media, when most cells have progressed into meiosis II, a time period after which *SOD1^LUTI^* expression normally ceases in cells synchronized by traditional nutritional cues alone (rather than the aforementioned *pGAL-NDT80* system). As expected, ectopic *SOD1^LUTI^* expression in late meiosis effectively blocked the synthesis of new protein seen at late meiotic timepoints in the vehicle control (Figure 4C). Thus, the precise timing of transient *SOD1^LUTI^* production controls a temporary cessation of Sod1 protein synthesis in mid-meiosis and its resurgence following the meiotic divisions, as gamete packaging is occurring.

### Sod1 loss is proteasome-dependent

Unlike in mitosis, dilution of cell contents through growth and cell division does not occur in meiosis. This means that a decrease in protein abundance indicates protein degradation under these conditions, which we have previously shown to be pervasive during budding yeast meiosis (Eisenberg and Higdon et al., 2018). The loss of Sod1 protein that we observed was dependent on the meiotic program, as cells lacking Ime1, the transcription factor required for the expression of early meiotic genes and meiotic entry, showed an increase in Sod1 protein levels relative to wild-type controls in matched sporulation media (Figure 5A). Furthermore, even in cells lacking LUTI-mediated *SOD1^canon^* downregulation, decreased Sod1 protein levels were observed in mid-meiosis (Figure 4B). The increase in Sod1 the absence of Ime1, and the decrease in its presence, began shortly after transfer to sporulation media, prior to the time that cells enter the meiotic divisions (Figure 5A). This early onset of degradation is consistent with the finding that cells arrested in late prophase due to lack of Ndt80 displayed normal robust degradation of Sod1 (Figure 5B). Altogether, these findings show that degradation of Sod1 begins early in the meiotic program.

**Figure 5.**
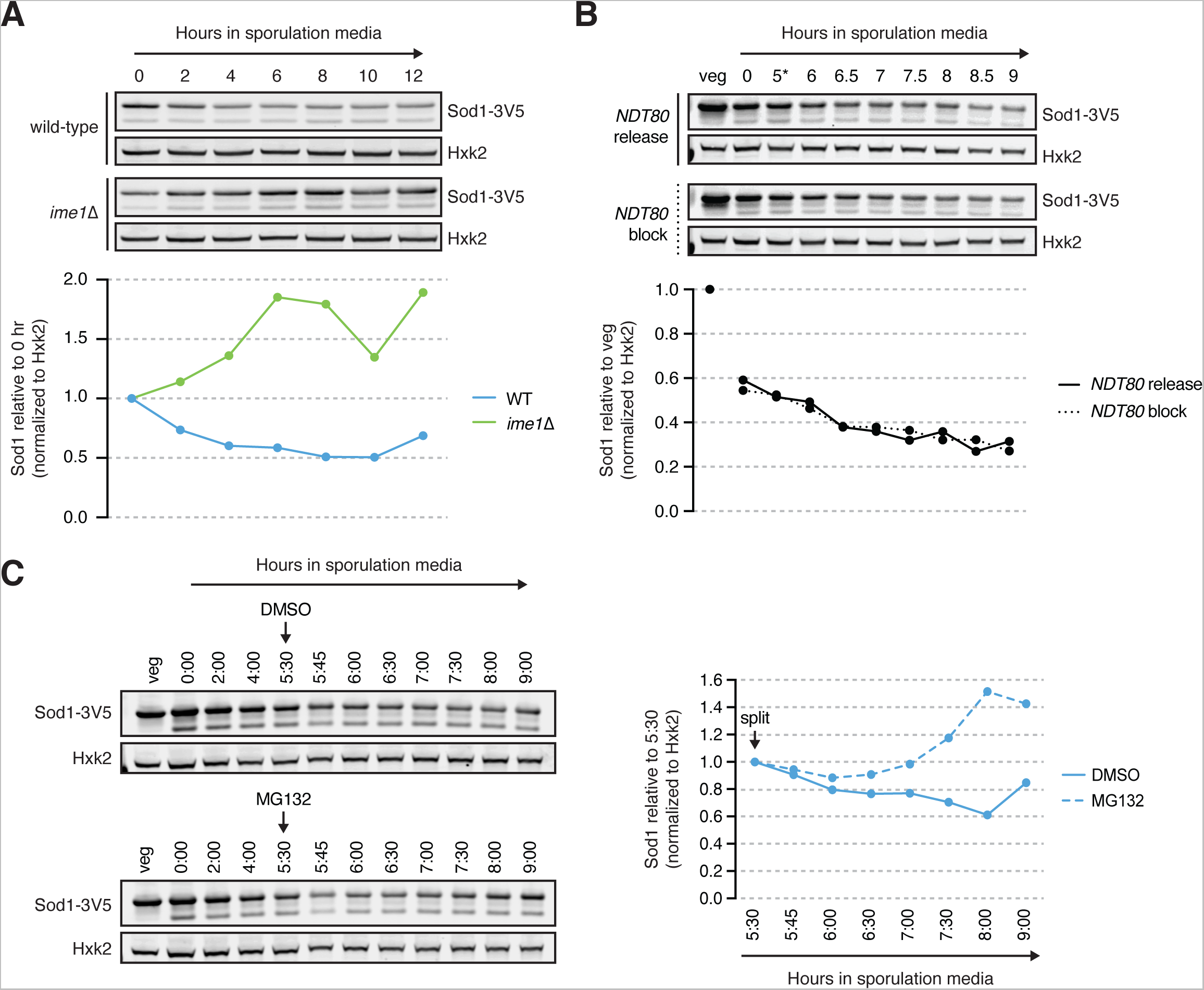
Sod1 loss is proteasome-dependent. (A) SDS-PAGE and immunoblotting for Sod1 in wild-type and *ime1*Δ cells throughout a meiotic time course (quantification shown below). (B) SDS-PAGE and immunoblotting for Sod1 in the presence or absence of *NDT80* expression (\**pGAL*-*NDT80* release at 5 hours; quantification shown below). (C) SDS-PAGE and immunoblotting for Sod1 in the presence or absence of the proteasome inhibitor MG132 (quantification shown to the right). Cultures were split after 5.5 hours in sporulation media and treated with either DMSO (vehicle control) or 100 µM MG132.

The ubiquitin proteasome system targets proteins for degradation by the proteasome, a large, ATP-powered complex. During budding yeast meiosis, virtually all components of the proteasome are upregulated. This upregulation peaks in late prophase at 5-fold above what is observed in mitotic cells (Brar et al., 2012; Cheng and Otto et al., 2018; Eisenberg and Higdon et al., 2018), which suggests elevated general proteasome activity. To determine if the meiotic decrease in Sod1 is mediated by the proteasome, we measured Sod1 levels in cells treated with the proteasome inhibitor MG132 compared to vehicle control-treated cells. Because the proteasome is essential for the meiotic divisions, we timed MG132 treatment to avoid disrupting the divisions but still within the window when Sod1 levels are decreasing. Upon MG132 treatment, after 5.5 hours in sporulation media, Sod1 protein levels did not continue to decline (Figure 5C), demonstrating that proteasome activity contributes to the mid-meiotic decrease in its abundance.

### Pre-meiotic Sod1 aggregates naturally occur and are cleared during the meiotic program

ALS-associated, aggregation-prone mutant hSOD1 exhibits increased turnover (Farrawell and Yerbury, 2021), leading us to consider the possibility that meiotic degradation of wild-type yeast Sod1 could be triggered by a change in its oligomerization status. To look for evidence of Sod1 aggregation, we performed immunofluorescence (IF) on fixed meiotic cells expressing endogenous Sod1-3V5. We found that over 90% of cells contained at least one bright focus at the time that they were resuspended in sporulation media (Figures 6A, 6B), consistent with the presence of Sod1 aggregates (Zeineddine et al., 2015; Gill et al., 2019). In contrast, mitotic cells in rich media did not contain bright foci of this nature and instead demonstrated heterogeneous, grainy Sod1 staining (YPD; Figure 6C). As cells progressed through meiosis, we saw disappearance of these foci with timing similar to the decrease in overall Sod1 protein levels, as assessed by SDS-PAGE (Figures 4A, 6A, 6B). Staging of cells fixed after 8 hours in sporulation media by DAPI staining revealed that only ∼10% of tetranucleate cells contain moderate foci, and bright foci were extremely rare, providing additional evidence that loss of foci takes place during the meiotic divisions (Figure 6B). Cells that were unable to enter the meiotic program due to the lack of Ime1 still formed Sod1 foci equivalently to matched wild-type controls, indicating that the trigger for focus formation is independent of meiotic entry (Figure 6D). However, the disappearance of Sod1 foci did not occur normally in cells lacking Ime1, which indicates that their removal is a programmed part of meiosis (Figure 6D).

**Figure 6.**
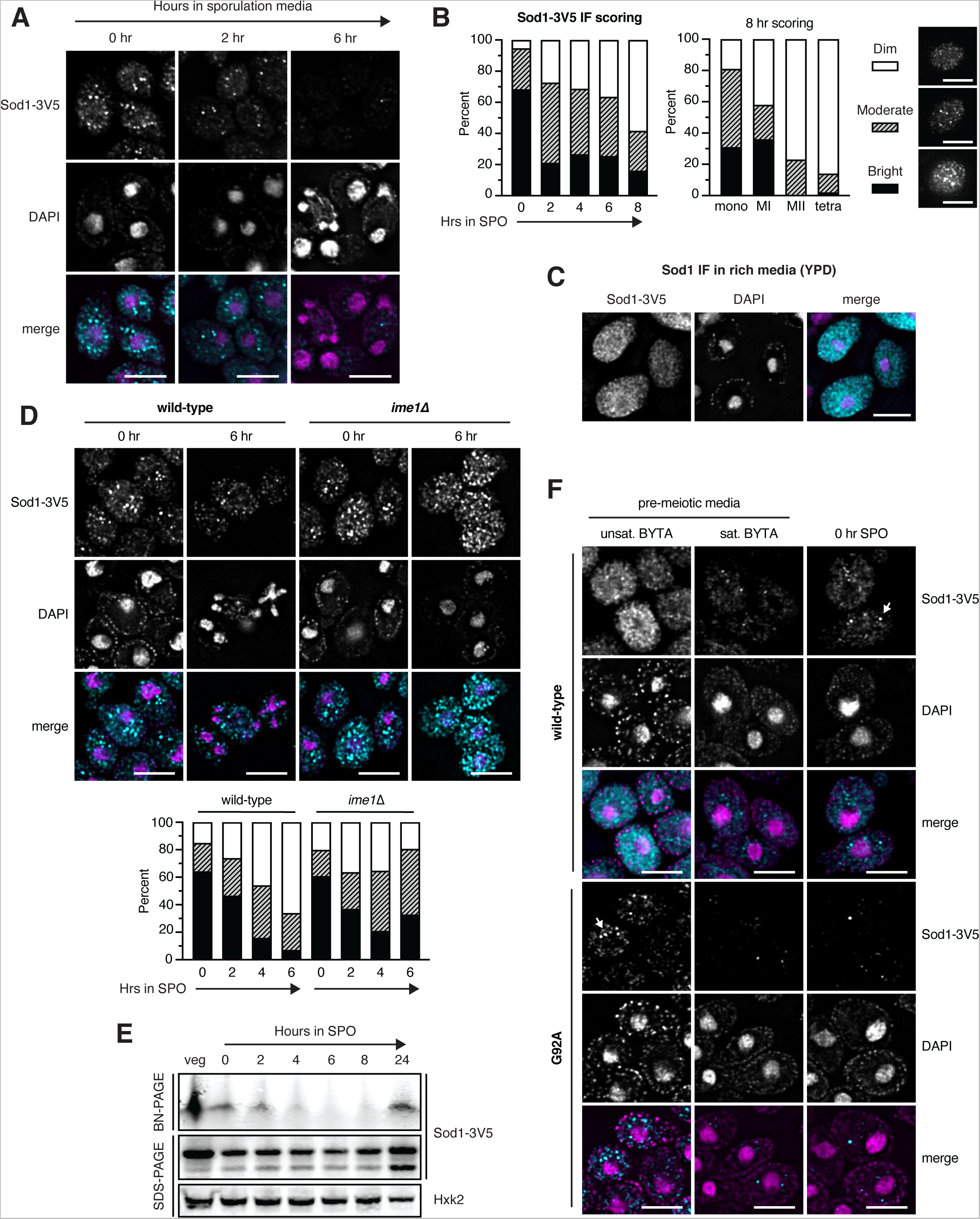
Pre-meiotic Sod1 aggregates naturally occur and are cleared during the meiotic program. (A) Immunofluorescence (IF) staining for Sod1 in cells fixed after 0, 2, and 6 hours in sporulation media. (B) Quantification of IF signal appearance in individual cells after 0-8 hours in sporulation media (left) and quantification of IF signal appearance in mononucleate (mono), meiosis I (MI), meiosis II (MII), and tetranucleate (tetra) cells from 8-hour samples (right). Scoring key shows representative examples of cells containing dim, moderate, or bright foci. At least 120 cells were scored per time point. (C) IF staining for wild-type Sod1 in rich media (YPD) during exponential growth. (D) IF staining for Sod1 in wild-type and *ime1*Δ cells after 0 and 6 hours in sporulation media. Quantification of Sod1 IF foci in wild-type and *ime1*Δ cells fixed after 0-6 hours in sporulation media is shown below. At least 150 cells were scored per time point. (E) Blue native PAGE (BN-PAGE) and immunoblotting for Sod1 during a meiotic time course. (F) IF staining for wild-type and G92A mutant Sod1-3V5 in pre-meiotic (unsaturated and saturated BYTA) and meiotic media (0 hr SPO). Identical exposure conditions were used during image acquisition, but post-acquisition exposures are different for wild-type and G92A micrographs to improve the visibility of G92A protein. All scale bars = 5 µm.

Properly folded and enzymatically active Sod1 exists as a 32 kDa homodimer. If Sod1 foci represented higher order multimers of Sod1, we would expect loss of this dimeric state by native gel analysis. Blue native polyacrylamide gel electrophoresis (BN-PAGE) and immunoblotting of endogenous Sod1-3V5 revealed abundant homodimer in vegetative native lysate (Figure 6E), but little to no dimeric Sod1 during mid-meiosis, in contrast to the relatively moderate dip in denatured protein observed by SDS-PAGE (Figures 4A, 6E, Figure S4A). By 24 hours in sporulation media, dimeric Sod1 becomes visible again, which we hypothesize represents new Sod1 synthesized late in sporulation. Because vacuolar protease activity is highly elevated during meiosis (Zubenko and Jones, 1981), we wondered if the disappearance of dimeric Sod1 might be due to degradation following cell lysis during sample preparation. However, preincubating vegetative lysate in meiotic lysate on ice for 30 minutes prior to performing BN-PAGE revealed a similar level of soluble, dimeric Sod1 as without preincubation (Figure S4B), arguing that post-lysis degradation by proteases is not likely to be causing the disappearance of Sod1 dimer in meiotic lysates. Further, overexpression of 3V5-tagged Sod1 in meiosis through addition of a *pATG8* transgene in strains housing endogenous, untagged Sod1 revealed soluble dimer during these mid-meiotic timepoints (Figure S4C), arguing that cellular conditions at these timepoints do not preclude the presence of soluble Sod1 dimer. We could not observe higher molecular weight species by BN-PAGE, which could be a result of the multiple centrifugation steps involved in the native extract preparation protocol. Together, these experiments support the model that formation of multimeric forms of wild-type Sod1 are triggered by pre-meiotic conditions, and that their removal is a natural aspect of the meiotic program.

Based on the focus formation observed by IF and the disappearance of soluble Sod1 dimer by BN-PAGE, we hypothesized that aggregation of Sod1 dimers was occurring prior to meiotic entry. If the foci we observed by IF indeed represented Sod1 aggregates, we hypothesized that their formation should be enhanced in cells carrying known aggregation-prone versions of Sod1. The G93A substitution was the first mutation that was well-characterized in mouse models of ALS and has since been widely studied in both mouse and human cell line research of the disease (Mejzini et al., 2019). To study the behavior of aggregation-prone Sod1 in meiosis, we introduced this mutation (G92A in yeast) within the endogenous *SOD1* gene. We performed IF on cells carrying either WT or G92A Sod1 in a variety of growth conditions, including exponential (unsat. BYTA) and saturated (sat. BYTA) growth in pre-meiotic media. These conditions represent the set of nutrient states used to grow cells prior to synchronous induction of meiosis upon transfer to sporulation media. When Sod1^WT^ cells were incubated in pre-meiotic media (BYTA), which replaces glucose with acetate as a carbon source, discrete bright foci emerged as the cultures became saturated (Figure 6F). In contrast, Sod1^G92A^ cells showed clear, bright foci even in unsaturated BYTA, indicating a higher propensity to aggregate than WT protein. This finding is consistent with the hypothesis that the Sod1 foci that we observed by IF represent aggregated Sod1.

### ALS-associated mutant Sod1 is rapidly degraded in meiosis

By IF, Sod1^G92A^ foci were readily observed in rich media and unsaturated growth in pre-meiotic media (Figure 6F). We noticed that these foci and almost all signal for Sod1^G92A^ were dramatically reduced in saturated BYTA, a condition in which cells begin to express Ime1, the transcription factor responsible for initiating the meiotic program. Based on this observation, and the correlation between the timing of wild-type aggregate disappearance and degradation that we observed, we hypothesized that aggregation of Sod1 prior to meiosis leads to clearance of these aggregates during meiosis. Consistent with this hypothesis and our IF data, Sod1^G92A^ protein levels dropped markedly and prematurely relative to Sod1^WT^ in the transition from rich media to sporulation media (Figure 7A). Similar results were observed by analysis of a more common ALS-associated and aggregation prone Sod1 mutant, A4V (A3V in yeast). After 24 hours in sporulation media, when wild-type protein levels have been restored, negligible amounts of Sod1^G92A^ and Sod1^A3V^ protein were present, indicating hyper-degradation, and decreased spore viability and poor colony growth were observed (Figure S5A).

**Figure 7.**
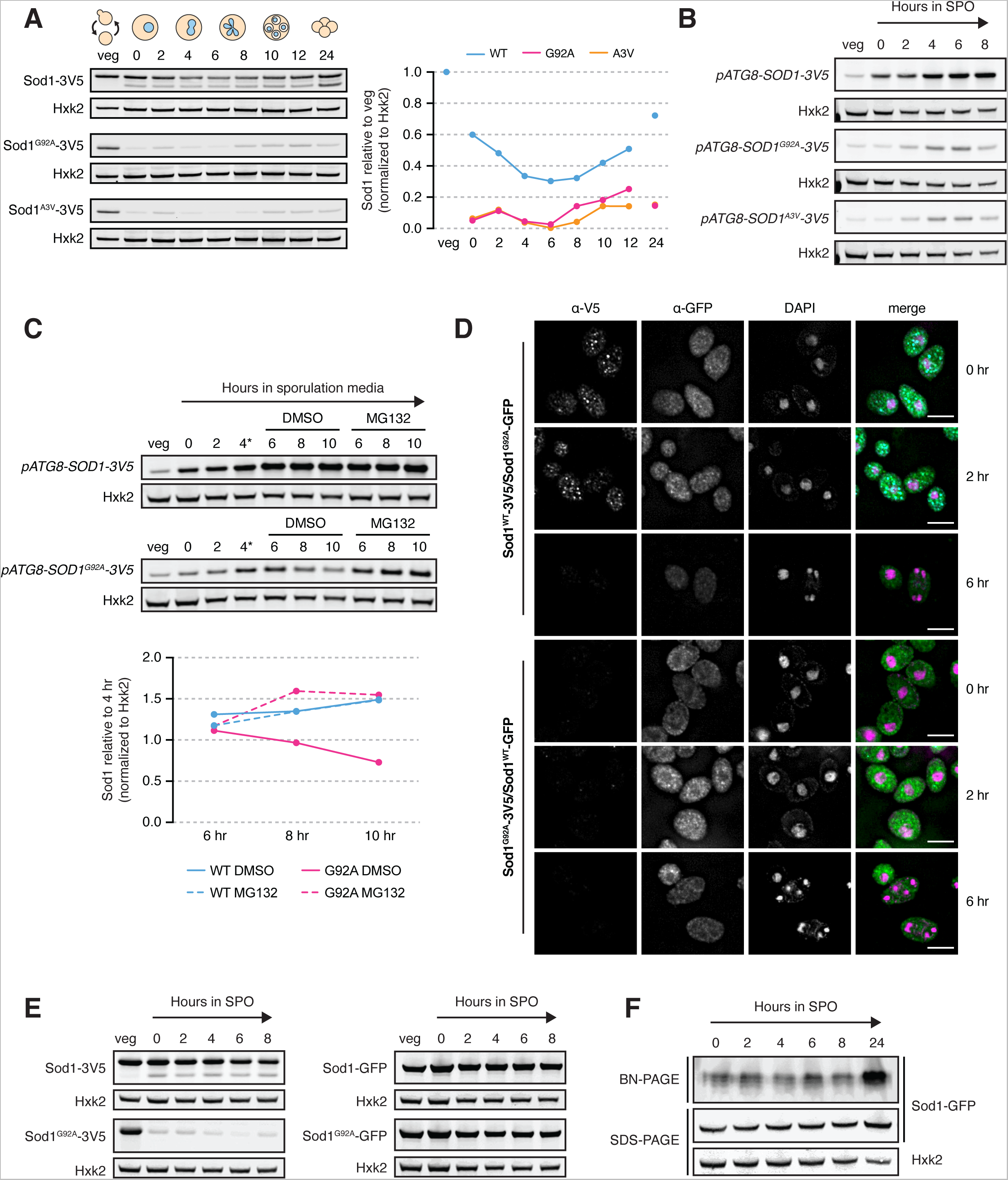
ALS-associated mutant Sod1 is rapidly degraded in meiosis. (A) SDS-PAGE and immunoblotting for Sod1^WT^, Sod1^G92A^, and Sod1^A3V^ throughout a meiotic time course (quantification shown to the right). (B) SDS-PAGE and immunoblotting for Sod1 during meiosis in strains expressing *pATG8*-driven transgenes. (C) SDS-PAGE and immunoblotting for Sod1 during meiosis in strains expressing *pATG8-SOD1^WT^-3V5* or *pATG8-SOD1^G92A^-3V5* (*after 4 hours in sporulation media, cultures were split and treated with either DMSO (vehicle control) or 100 µM MG132, quantification shown below). (D) IF staining for Sod1^WT^ or Sod1^G92A^ tagged with either 3V5 or GFP in trans-heterozygous strains after 0, 2, and 6 hours in sporulation media. (E) SDS-PAGE and immunoblotting for Sod1^WT^ or Sod1^G92A^ tagged with either 3V5 or GFP in vegetative and meiotic media conditions. (F) Blue native PAGE (BN-PAGE) and immunoblotting for Sod1-GFP during a meiotic time course. All scale bars = 5 µm.

We aimed to probe whether the increased degradation of aggregation-prone mutant versions of Sod1 occurred by the same proteasome-mediated route that we found to decrease wild-type Sod1 levels in meiosis, but inhibition of the proteasome prior to meiotic entry prevents cells from entering meiosis, when the bulk of degradation in these mutants was seen (Figure 7A). Thus, we designed a strategy that would allow us to express Sod1^G92A^ and Sod1^A3V^ protein at high levels in meiosis, expressing either wild-type or ALS mutant Sod1-3V5 from an ectopic locus under the control of a strong promoter (*pATG8*) that drives especially high expression in mid- to late-meiosis. SDS-PAGE analysis showed that Sod1^G92A^ levels were substantially lower than wild-type protein expressed from the *ATG8* promoter (Figure 7B), despite no appreciable differences in transcript abundance by RT-qPCR (Figure S5B). We observed similar results using cells carrying *pATG8-Sod1^A3V^-3V5*, but not those carrying human wild-type SOD1 (Figure S5C). Given the comparable mRNA levels and our previous finding that Sod1 is degraded by the proteasome during meiosis (Figure 5C), we next split cultures carrying *pATG8*-driven Sod1^WT^ and Sod1^G92A^ after four hours in sporulation media and treated with either DMSO or MG132 to inhibit the proteasome. Monitoring protein expression revealed elevated, although not fully rescued, levels of meiotic expression of the overexpressed mutant protein with MG132 treatment (Figure 7C), indicating that the reduced expression levels of the ALS mutant Sod1 protein relative to wild-type protein are due to proteasome-mediated degradation.

To this point, our data support the model that pre-meiotic conditions in yeast drive aggregation of wild-type Sod1 protein and the meiotic program drives removal of aggregates and degradation of Sod1. The timing of aggregate disappearance and the efficient degradation of aggregation prone ALS-associated mutant versions of Sod1 suggested that aggregates are targeted for degradation. This model would predict that a version of Sod1 that does not aggregate should not be degraded during the meiotic divisions. To test this, we analyzed cells with endogenously GFP-tagged Sod1. Previous work with Sod1-GFP has yielded confusing results, as it does not seem capable of forming aggregates, even when mutated at the sites representing aggregate-prone ALS mutations (Bastow et al., 2016). Consistent with published studies, we observed a relatively even distribution of endogenous Sod1^WT^-GFP and Sod1^G92A^-GFP in fixed cells in early meiosis (Figure S5D). Immunofluorescence also revealed no Sod1 foci for GFP-tagged protein, even when foci for Sod1-V5 were seen in the same diploid cells carrying one endogenous copy of Sod1 tagged with GFP and one with 3V5 (Figure 7D). Consistent with our microscopy data, SDS-PAGE analysis showed that both wild-type and G92A Sod1 were much more stable throughout meiosis when tagged with GFP, in contrast to the pattern seen for V5 or untagged Sod1 (Figures 7E, 4A, 1A), as expected if preventing aggregation of Sod1 also prevents its meiotic degradation. Furthermore, BN-PAGE analysis of Sod1-GFP revealed soluble dimer throughout meiosis (Figure 7F) and Sod1^G92A^-GFP rescued the reduced spore viability and poor colony growth seen in cells expressing Sod1^G92A^-3V5 (Figure S5E). These data support the model that a failure of Sod1 to aggregate prevents its meiotic degradation.

Together, our findings show that aggregation of existing wild-type Sod1 protein naturally occurs in pre-meiotic conditions, and that clearance of these aggregates is a programmed aspect of meiotic differentiation that begins early in the meiotic program. As clearance continues, Sod1 protein synthesis is halted during the meiotic divisions by the transient expression of a LUTI mRNA that is responsive to the UPR^ER^. Loss of this repressive transcript allows Sod1 synthesis to resume, which we propose promotes the formation of healthy, viable gametes through repopulation of the Sod1 pool with new protein (Figure 8).

**Figure 8.**
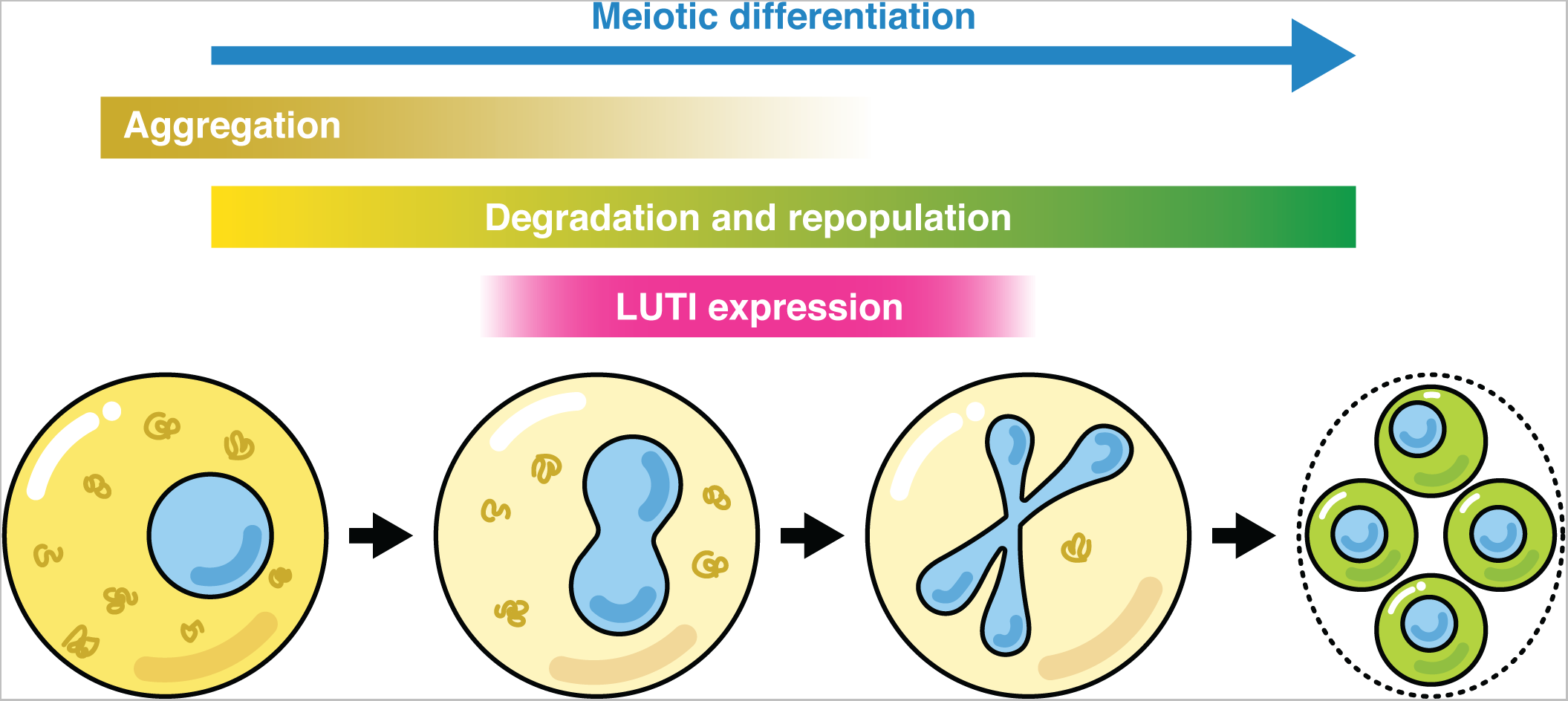
Model for the meiotic regulation of Sod1 protein during budding yeast meiosis. Prior to entry into the meiotic program, a population of Sod1 is present in an aggregated form. When cells begin the meiotic differentiation program, upregulation of degradation factors, including the ubiquitin-proteasome system, leads to degradation of preexisting Sod protein. During the meiotic divisions, a gradual disappearance of these Sod1 aggregates occurs, which coincides with the expression of the UPR^ER^-driven *SOD1^LUTI^* mRNA isoform that acts to antagonize *SOD1^canon^*. expression and therefore the synthesis of new Sod1 protein. LUTI expression ceases around the time that most cells no longer contain observable Sod1 aggregates, and restoration of canonical mRNA expression allows for the repopulation of cells with new Sod1 protein, which we hypothesize to be important for the generation of healthy, viable gametes.

## DISCUSSION

The primary goal of gametogenesis is packaging of the full set of cellular materials required to produce a highly fit next generation. This includes creation of genetic diversity and segregation of critical cellular components. It also involves clearance of cellular damage, resulting in gametes that act as “young” cells, regardless of progenitor cell age (Ünal et al., 2011). The factors that contribute to this natural rejuvenation are not yet known, but removal of preexisting nuclear, mitochondrial, and ER proteins has recently been shown to accompany the meiotic program (Sawyer et al., 2019; King and Goodman et al., 2019; Otto et al., 2021). It is intriguing that, more broadly, many abundant “housekeeping” proteins are degraded during the meiotic program in budding yeast and resynthesized prior to gamete maturation (Eisenberg and Higdon et al., 2018). These proteins, which include ribosomal subunits, nuclear pore complex components, and Sod1, are not typically degraded during mitotic division. However, their quality is critical to core cellular functions, and it is possible that ensuring this quality motivates the high energetic cost of their clearance and resynthesis. Here, we identify the complex set of regulatory events that enable resetting of Sod1 levels, an abundant and primarily cytosolic protein that is critical for combatting oxidant-based cellular damage.

The appearance of aggregates of wild-type Sod1 in pre-meiotic cells was surprising and is, to our knowledge, the first time this has been observed to occur pervasively in healthy, wild-type cells. Why do they form in pre-meiotic conditions? This remains an open question, but pre-meiotic media leads to increased respiration, which causes oxidative damage (Semchyshyn et al., 2011). In stationary-phase yeast, His71, His120, and Cys146 are oxidized, which results in the generation of enzymatically inactive, soluble protein aggregates (Martins and English 2014). However, an in-gel activity assay showed that Sod1 remains enzymatically active throughout meiosis, even in an early meiotic time point when foci are most abundant by immunofluorescence (Figure S4D). Because Sod1 is only catalytically active as a dimer, we believe that the foci we observe represent clusters of dimeric and catalytically active Sod1 that are SDS-soluble. The nature of Sod1 aggregates observed upon mutation or in disease states has been controversial, with both SDS-soluble large aggregates and SDS-insoluble amyloids reported (Brotherton et al., 2013; Karch et al., 2009; Basso et al., 2009). It remains unclear the degree to which the foci that we observe resemble species seen in human pathogenic states, and defining their physical nature is an important future direction.

It is interesting to note that other protein aggregates have been shown to be cleared during gametogenesis in yeast. These include Hsp104-bound aggregates, which are thought to represent misfolded proteins, and which are prominent in aged cells (Ünal et al., 2011), as well as the natural Rim4 amyloid, which regulates translation of several key meiotic mRNAs (Berchowitz et al., 2015). The removal of Sod1 foci begins before clearance of both of these aggregate classes, indicating that multiple parallel routes to aggregate removal are part of the meiotic program.

What is the purpose of pre-meiotic Sod1 aggregation? It may simply be a side-effect of the nutritional starvation that accompanies pre-meiotic conditions, or it may occur for the purpose of enabling meiotic degradation of preexisting Sod1 protein. Several lines of evidence suggest that, regardless of why aggregation occurs, it does facilitate Sod1 clearance. This evidence includes the timing of Sod1 foci disappearance which corresponds with the drop in total Sod1 protein levels, the hyper-degradation observed in Sod1 mutants that prematurely aggregate in pre-meiotic media, and the lack of degradation observed in cells expressing Sod1-GFP, which do not show evidence of aggregation. Although we were surprised to find such a striking difference between the aggregation and meiotic degradation of 3V5- vs. GFP-tagged Sod1, there is precedent for GFP-tagging of Sod1 altering its aggregation behavior *in vitro* (Stevens et al., 2010). Identifying the specific mechanism of Sod1 degradation in meiosis will clarify whether the aggregated and/or soluble Sod1 pools are targeted. At least a subset of meiotic Sod1 degradation is dependent on the proteasome, but the molecular adaptors involved remain unidentified, and it is also possible that autophagy also acts parallel to remove meiotic Sod1. The failure of proteasome inhibition to fully rescue expression of Sod1^G92A^ driven by the *ATG8* promoter is consistent with this possibility (Figure 7C). Nevertheless, the active degradation of this enzyme in meiosis contrasts with its regulation in nearly all other studied contexts.

Sod1 is required for gamete viability in budding yeast (Figure 1C), but its specific critical function in gametogenesis is not known. Given the high degree of respiration required for meiosis, downregulation of Sod1 protein levels during this process may seem counterintuitive, but recent work argues that although it is best known for its antioxidant role, only a small fraction of this abundant enzyme is needed to protect cells from superoxide (Montllor-Albalate et al., 2019). This discrepancy was puzzling until Sod1 was found to play additional roles, including nutrient sensing and redox homeostasis (Reddi and Culotta, 2013; Montllor-Albalate et al., 2022). Identifying the specific roles that underlie its importance for meiosis is an interesting future area of research.

The cessation of Sod1 synthesis and its replenishment late in meiosis are driven by the activation and abatement, respectively, of *SOD1^LUTI^* expression. We found that this timing mirrors UPR^ER^ activation timing in meiosis, which we also found to be capable of *SOD1^LUTI^* production in a mitotic context. Our previous mRNA-seq study of cells treated with DTT did not identify *SOD1^LUTI^* as a Hac1 target (Van Dalfsen et al., 2018). However, this study used a highly stringent cut-off for UPR^ER^-dependent LUTI production and the level of *SOD1^LUTI^* observed in this current study in response to mitotic UPR^ER^ activation is modest compared to the targets previously identified, although prominent in meiosis. This, together with the fact that *SOD1 ^LUTI^* production is dependent on the UPR^ER^ but not UPREs suggests that either Hac1 is increasing the expression or activity of an unknown factor that is subsequently turning on *SOD1^LUTI^* or that Hac1 binds another sequence within the LUTI promoter. Given that the promoters of many validated Hac1 targets do not contain predicted UPREs (Travers and Patil et al., 2000; Patil et al., 2004), investigation of new modes of Hac1-induced transcription is warranted.

The LUTI isoform of *SOD1* is nearly four times the length of *SOD1^canon^*., which is to our knowledge the most substantial size difference between LUTI and canonical isoforms observed to date and is especially remarkable when considering the compact nature of the *S. cerevisiae* genome. The ability of LUTI-based regulation to temporarily pause production of housekeeping genes that are “on” by default is interesting. Such genes may be activated by multiple transcription factors and thus timed repression may be best achieved by simply silencing all transcription from the canonical transcription start site with a LUTI rather than use of specific transcriptional repressors. In the case of *SOD1,* timed toggling between two transcript isoforms of differential translatability, together with regulated aggregation and degradation, act in concert to remove a subset of preexisting Sod1 protein from eligibility for gamete inheritance and simultaneously replenish the Sod1 pool with new protein (Figure 8). This elegant set of regulatory mechanisms achieves the regeneration of a housekeeping protein with established links to prevention of cellular aging. It is thus intriguing that it occurs during gametogenesis, a natural context of cellular rejuvenation. “Housekeeping” protein replenishment in gametes represents an exciting theme, and is also seen for the many other proteins, including other antioxidant enzymes and the proteins that make up ribosomes (Eisenberg and Higdon et al., 2018).

Beyond its potential significance to gametogenesis, our findings also point to a new natural context in which to study important and unresolved aspects of Sod1’s role in human disease. Although SOD1 abnormalities are most famously associated with ALS, its dysregulation is also seen in Down syndrome (Engidawork and Lubec, 2001), Parkinson’s (Trist et al., 2017), and cancer (Papa et al., 2014; Tsang et al., 2018; Liu et al., 2020; Wang et al., 2021). Sod1 aggregates have been proposed to lead to cellular toxicity, although it has also been argued that they serve a protective role in isolating populations of damaged hSOD1 to be targeted for degradation (Zhu et al., 2018; Gill et al., 2019). Discriminating between these hypotheses is not possible based on clinical samples and is challenging in complex mammalian model systems. The natural aggregation and clearance of Sod1 that we observe to occur as part of the meiotic program in budding yeast suggest that future studies of Sod1 in yeast meiosis have the potential to improve our understanding of how this protein is regulated in a growing number of human diseases and conditions.

## MATERIALS AND METHODS

### Yeast strains and plasmids

All strains used in this study are derived from the SK1 strain of *Saccharomyces cerevisiae*. Strain genotypes are listed in Supplementary file 1. The following alleles were constructed in previous studies: *pGAL-NDT80 GAL4-ER* (Carlile and Amon, 2008), lexA-B112-ER (Chia et al., 2017), *hac1*Δ (Van Dalfsen et al., 2018), *pdr5*Δ (Sawyer et al., 2019).

To visualize Sod1 for immunoblotting and immunofluorescence, we C-terminally tagged endogenous Sod1 with 3V5 or yeast codon-optimized EGFP (yoEGFP). To induce *SOD1^LUTI^* mRNA overexpression, we engineered a strain expressing a chimeric lexA-B112 transcription factor with an estradiol-binding domain (lexA-B112-ER) and 8 tandem lex operator sequences (8lexO) 1680 bp upstream of the *SOD1* ORF ATG. Positioning of the 8lexO array was informed by mRNA sequencing (Brar et al., 2012) and transcript leader sequencing (TL-seq) data (Chia et al., 2021). To overexpress either wild-type, G92A, A3V, or yeast codon-optimized human Sod1 in meiosis, we generated strains with *pATG8*-driven transgenes integrated at the HIS3 locus. To delete the *SOD1* gene, the ORF was replaced with a KanMX6 marker in a wild-type, diploid strain. Transformant diploids were sporulated, then *sod1*Δ haploids were isolated and mated to generate a homozygous *sod1*Δ diploid strain.

LUTI disruption (*CYC1t* insertion), UPRE mutations, G92A and A3V mutations, and unmarked C-terminal 3V5 Sod1 tagging were performed using CRISPR-Cas9 (Jinek et al., 2012; Anand et al., 2017). To disrupt *SOD1^LUTI^* expression, the termination sequence from the *CYC1* gene (*CYC1t*) was inserted between the LUTI and canonical TSSs. The 248 bp *CYC1* terminator sequence was amplified from plasmid pÜB196 and inserted into the genome 278 bp upstream of the *SOD1* ORF ATG. To mutate the unfolded protein response elements (UPREs) upstream of the LUTI TSS, the proximal UPRE (5′-ACACGT-3′) was deleted and the distal UPRE was scrambled (5′-TCGTGG-3′ to 5′-CCTAGG-3′). To target Cas9 to specific sites within the genome, Golden Gate cloning (Engler et al., 2008) was used to insert 20-nucleotide guide RNA (gRNA) sequences (found in Supplementary file 4) into a centromeric plasmid containing a URA3 marker and pPGK1-Cas9 (a gift from Gavin Schlissel, University of California, Berkeley, CA). Plasmids were then co-transformed into yeast cells with a repair template (see Supplementary file 2 for details). All repair templates used in this study were designed to simultaneously mutate the PAM sequence adjacent to the gRNA sequence. Once strains were sequence-validated, cells were cured of plasmid expression by streaking for singles on YPD and selecting colonies that no longer exhibited growth on –URA.

### Yeast growth and sporulation conditions

To prepare cells for sporulation, diploid strains were patched out onto YPG (2% glycerol) plates and incubated at 30°C overnight. In the morning, strains were patched onto YPD 4% plates (1X YEP with 4% dextrose) and incubated at 30°C. Later that day, YPD (1X YEP with 2% dextrose, supplemented with Trp and Ura) cultures were inoculated with cells from YPD 4% plates and incubated at room temperature for 24 hours. Once YPD culture volumes had reached a minimum OD_600_ of 10, cells were used to inoculate BYTA (2% bacto tryptone, 1% yeast extract, 1% potassium acetate, 50 mM potassium phthalate) cultures at a starting OD_600_ of 0.25 and then incubated at 30°C overnight. Once cultures reached a minimum OD_600_ of 5, cells were pelleted at 1,900 rcf for 2 minutes, washed once with autoclaved MilliQ water, and resuspended to an OD_600_ of 1.85 in sporulation (SPO) media (2% (w/v) potassium acetate (pH 7), 0.02% (w/v) raffinose, supplemented with adenine, uracil, histidine, leucine, and tryptophan) and incubated at 30°C. To maintain proper aeration, all YPD, BYTA, and SPO culture volumes prepared for meiotic experiments did not exceed 1/10^th^ of flask volumes.

For certain experiments, meiotic cultures were synchronized using the *pGAL-NDT80 GAL4-ER* system (Carlile and Amon 2008). After incubation at 30°C for 5 hours, meiotic cultures were released from prophase I arrest by inducing *NDT80* expression with the addition of β-estradiol (in 100% ethanol) to achieve a final concentration of 1 µM. For *SOD1^LUTI^* overexpression in mitosis or meiosis, expression was induced by adding β-estradiol to a final concentration of 30 nM. For vehicle control samples, an equal volume of 100% ethanol was added.

For vegetative experiments, strains were patched out onto YPG plates and incubated at 30°C overnight. The following morning, strains were patched onto YPD plates (1X YEP with 2% dextrose for haploids or 4% dextrose for diploids) and incubated at 30°C. Later that day, YPD cultures were inoculated with cells from YPD plates and incubated at 30°C overnight. The next day, YPD cultures were backdiluted to an OD_600_ of 0.05. For both LUTI overexpression and UPR^ER^ experiments, backdiluted YPDs were incubated at 30°C for 4 hours, split into two cultures, and treated with either a drug or vehicle control, and incubated at 30°C for 1-2 hours. For matched vegetative samples accompanying meiosis experiments, backdilutions were prepared from the same saturated YPD cultures used to inoculate BYTAs and were incubated at 30°C for 4-5 hours prior to sample collection.

### UPR^ER^ induction

To activate the UPR^ER^ during vegetative exponential growth, cells were treated with stock solutions of dithiothreitol (DTT) or tunicamycin. For each experiment, a 1M stock solution of DTT (cat. no. 15508-013, Invitrogen) was prepared fresh, then added to cultures to achieve a final concentration of 5 mM. For tunicamycin (cat. no. 654380, Calbiochem) treatment, 25 mg/mL stocks (prepared in DMSO) were thawed from -80°C and added to cultures to achieve a final concentration of 2 µg/mL.

### Proteasome inhibition

To inhibit the proteasome, cell cultures were treated with a 100 mM stock of MG132 (Z-Leu-Leu-Leu-al, C2211, Sigma) in DMSO to achieve a final concentration of 100 µM. All strains used in MG132 experiments have the *PDR5* gene deleted, which encodes an ABC transporter, allowing for efficient uptake and retention of the drug (see strain genotypes listed in Supplementary file 1).

### Spore viability

To assess spore viability, 100 µL aliquots of SPO cultures collected after 24 hours at 30°C were spun in 1.5 mL tubes at 1,900 rcf for 2 minutes at room temperature and SPO was removed from cell pellets by pipetting. To digest asci and allow for manual separation of individual spores on agar plates, cells were resuspended in 20 µL of [1 mg/mL] 100T zymolyase (cat. no. 08320932, MP Biomedicals or cat. no. 320932, VWR) and incubated at room temperature for 6-7 minutes. To stop digestion, 180 µL of MilliQ H_2_O was added to each sample, and 20 µL of digested cells were then pipetted down the midline of YPD agar plates (1X YEP with 2% dextrose). Isolation of spores from the same tetrad was performed by microdissection using a fiberoptic needle attached to a Zeiss light microscope. For each experiment, 10-20 tetrads were dissected per genotype.

### RNA extraction

To extract total RNA, 3.7-18.5 OD_600_ units of cells were spun at room temperature (1,900 rcf for vegetative samples and maximum speed for meiotic samples) for 1 minute. After removing the supernatant, cell pellets were frozen in liquid nitrogen and stored at -80°C. Pellets were thawed on ice and resuspended in 600 µL each of TES (10 mM Tris pH 7.5, 10 mM EDTA, 0.5% SDS) buffer and acid phenol (Sigma P4682) and shaken at 1,400 rpm in a Thermomixer^®^ R (Eppendorf) for one hour at 65°C. Samples were incubated on ice for 5 minutes, then spun at maximum speed for 10 minutes at 4°C. The aqueous phase was transferred to pre-chilled tubes containing 600 µL fresh acid phenol, samples were vortexed briefly to mix, then spun again at maximum speed for 10 minutes at 4°C. The aqueous phase was transferred to pre-chilled tubes containing 600 µL chloroform, vortexed briefly to mix, and spun at maximum speed for 5 minutes at 4°C. The aqueous phase was then transferred to pre-chilled 1.5 mL low-adhesion tubes containing 700 µL 100% isopropanol and 60 µL 3M NaOAc pH 5.2. Tubes were inverted to mix, then stored at -20°C for a minimum of 2 hours before pelleting at 13,000 rpm at 4°C for 5 minutes. After aspirating off the supernatant, RNA pellets were dried in a fume hood for 30-60 minutes until clear. Once dried, pellets were resuspended DEPC treated water (volume varied depending on pellet size, but usually ranged from 25-100 µL). RNA samples were stored at -80°C and were thawed on ice and quantified on a NanoDrop 2000 (Thermo Scientific) prior to northern blotting or cDNA synthesis.

### Northern blotting

RNA samples were thawed from -80°C on ice. Once thawed and quantified, 10 µg aliquots of RNA were speed-vacuumed (Savant Speed Vac^©^ SPD111V), resuspended in 1X NorthernMAX™ dye (Invitrogen, REF AM8552), and denatured at 65°C for 15 minutes. Samples were then cooled on ice before being loaded into formaldehyde gels (6.66% formaldehyde, 1.2% agarose) and separated by electrophoresis in 1X MOPS buffer (1X MOPS, 2 mM NaOAc, 1 mM EDTA, pH 7). RNA was then transferred to Hybond^®^ N^+^ membranes (cat. no. RPN203B, GE Healthcare) by capillary action using 10X SSC (1.5M NaCl, 150 mM Na_3_Citrat-2H_2_O) overnight. After transferring, RNA was UV-crosslinked to membranes using a crosslinker and rRNA was visualized by staining with methylene blue (0.3M NaOAc, 0.02% methylene blue, pH 5.5) for 5 minutes at room temperature. Excess methylene blue was removed by washing membranes with source MilliQ (5X quick washes followed by 3X 2-minute washes), then membranes were imaged on a Gel Doc XR+ (Bio-Rad). Membranes were then incubated with 22 mL ULTRAhyb™ hybridization buffer (cat. no. AM8669, Invitrogen) and 500 µL boiled sonicated salmon sperm DNA (cat. no. 201190-81, Agilent) for a minimum of 30 minutes at 68°C. Probe synthesis was performed as directed using the MaxiScript kit (cat. no. AM1314, Invitrogen). The *SOD1* ORF probe template was synthesized by amplifying wild-type SK1 genomic DNA using primers with the sequences 5′-CAACCACTGTCTCTTACGAGATCGC-3′ and 5′-taatacgactcactataggCACCATTTTCGTCCGTCTTTACG-3′ (lowercase letters denote T7 promoter), which amplifies 201 bp of the *SOD1* ORF. Upon addition of 2 µL T7 enzyme mix (REF 2719G, Invitrogen) and 5 µL 32α-UTP, probe mix was incubated at 37°C for 15 minutes. After synthesis, 1 µL of TURBO DNAse (REF 2238G2, Invitrogen) was added and probes were incubated at 37°C for 10 minutes. Before spinning the probe mix through a hydrated NucAway™ column (REF AM10070, Invitrogen), 1 µL of 0.5M EDTA pH 8 was added. Probes were eluted through the columns by spinning at 3,000 rpm for 3 minutes at room temperature, then added to hybridization tubes and incubated at 68°C overnight. Low stringency (2X SSC, 0.1% SDS) and high stringency (0.1X SSC, 0.1% SDS) wash buffers were heated to 68°C prior to use. After overnight hybridization, excess probe mix was poured off, and membranes were washed 2X with low stringency buffer (10 minutes per wash) and 3X with high stringency buffer (15 minutes per wash) to remove any unbound probe. Phosphor imaging screens (Molecular Dynamics) were then exposed to membranes overnight and RNA was visualized using a Typhoon scanner (Typhoon 9400, Molecular Dynamics).

### Single-molecule RNA fluorescence *in situ* hybridization (smRNA-FISH)

Single-molecule RNA FISH was performed as described in Chen et al., 2017 and Chen et al., 2018 (an adaptation of Raj et al., 2008 for budding yeast). Briefly, cells were fixed in 3% formaldehyde, incubated at room temperature for 20 minutes, then incubated at 4°C overnight. For meiotic experiments, cells were fixed after 6 hours in sporulation media. For mitotic experiments involving UPR^ER^ induction, cells were fixed one hour after drug treatment. The next day, cells were washed, digested, and hybridized with fluorescent probe sets in a formamide-based solution in the dark overnight. The probe sets hybridizing to the LUTI-specific sequences and sequences within the *SOD1* ORF were conjugated to fluorophores Quasar^®^ 670 and CAL Fluor^®^ 590, respectively (Stellaris), and probe sequences are listed in Supplementary file 2. The following day, cells were washed and incubated with DAPI and anti-bleach reagents prior to imaging on a DeltaVision Elite Microscope (GE Healthcare). For all experiments, DAPI was visualized by UV excitation (exposure and transmission varied between experiments depending on staining intensity), LUTI probes were visualized by CY-5 excitation (1.3s exposure, 100% transmission), ORF probes were visualized by TRITC excitation (1.3s exposure, 100% transmission), and reference images were taken mid-stack using polarized light (0.1s exposure, 32% transmission). Quantification of deconvolved smRNA-FISH images was performed using StarSearch (Raj et al., 2008) and cell boundaries were defined manually by tracing cells within a REF image using a drawing tablet (Wacom).

### RT-qPCR

cDNA was prepared by first strand synthesis using the TURBO DNA-*free*™ Kit (REF AM1907, Invitrogen) and Superscript III (REF 5675, Invitrogen). First, 2.5 µg of total RNA was treated with DNase I and incubated at 37°C for 30 minutes. After inactivation with DNase Inactivation Reagent, 2 µL of DNase-treated RNA was combined with 4.5 µL of a master mix containing random hexamers Roche and dNTPs. Samples were incubated at 65°C for 5 minutes, then placed on ice. While on ice, 3.5 µL of RT master mix (1X first-strand buffer, DTT, RNaseOUT, SSIII [REF 56575, Invitrogen]) was added to each tube, then samples were incubated at 25°C for 5 minutes, 50°C for 1 hour, and 70°C for 15 minutes. For RT-qPCR, 5.2 µL Absolute Blue SYBR Green ROX (REF AB-4162, Thermo Scientific) and 4.8 µL cDNA (diluted 1:20 prior to use) was added to each well of qPCR plates (REF 4346906, Life Technologies). Samples were run on a StepOnePlus™ RT-qPCR machine (Applied Biosystems). Average C_T_ values of triplicate reactions were used to calculate expression changes. RT-qPCR data was normalized to *ACT1* for mitotic experiments and *PFY1* for meiotic experiments (primers listed in Supplementary file 2).

### SDS-PAGE immunoblotting

To prepare lysates for SDS-PAGE, 3.7-9.25 OD_600_ units were pelleted at 1,900 rcf for 2 minutes at 4°C, resuspended in 2 mL of 5% trichloroacetic acid (SA433, Fisher) and incubated at 4°C for a minimum of 20 minutes. Samples were then spun at 1,900 rcf for 2 minutes at room temperature and cell pellets were washed by vortexing with 1 mL of 100% acetone. Samples were spun at maximum speed at room temperature for 5 minutes, acetone was pipetted off, and pellets were dried in a fume hood overnight. Once dried, pellets were resuspended in 100 µL of lysis buffer (50 mM Tris pH 7.5, 1 mM EDTA, 3 mM DTT, 1X cOmplete protease inhibitor cocktail [SKU, 11836145001, Roche], 1.1 mM PMSF, and 1X Pepstatin A) and beaten on a bead-beater for 5 minutes at room temperature with 100 µL of acid-washed glass beads (cat. no. G8772, Sigma). Next, 50 µL of 3X SDS buffer (18.75 mM Tris pH 6.8, 6% β-mercaptoethanol, 30% glycerol, 9% SDS, 0.05% bromophenol blue) was added, and samples were boiled at 95°C for 5 minutes. After boiling, samples were cooled at room temperature for at least 5 minutes before storing at -20°C.

Prior to loading, frozen SDS-PAGE samples were incubated at 37°C for 5 minutes, then spun at max speed for 5 minutes at room temperature. 4-8 µL of samples and 3 µL of Precision Plus Protein Dual Color Standard (cat. no. 1610374, Bio-Rad) were run on 4-12% Bis-Tris Bolt gels (Thermo Fisher) and transferred to 0.45 µm nitrocellulose membranes (Bio-Rad) with cold 1X Trans-Blot^®^ Turbo™ buffer in a semi-dry transfer apparatus (Trans-Blot^®^ Turbo™ Transfer System, Bio-Rad). Membranes were incubated at room temperature for 90 minutes in a 1:1 blocking solution of 1X PBS and PBS Odyssey^®^ Blocking Buffer (cat. no. 927-4000, LI-COR) or PBS Intercept^®^ Blocking Buffer (cat. no. 927-70001, LI-COR) and incubated in primary antibody solutions at 4°C overnight. Membranes were then washed 4X with 1X PBS-(0.01%)T (5 minutes per wash) at room temperature before incubating in secondary antibody solutions at room temperature for 150 minutes. Membranes were washed 3X with PBS- (0.01%)T and once with 1X PBS (5 minutes per wash) at room temperature prior to imaging with the Odyssey^®^ system (LI-COR Biosciences).

All primary and secondary antibodies were diluted in PBS Odyssey^®^ Buffer or PBS Intercept^®^ Buffer with 0.01% tween-20. Primary antibodies: mouse α-V5 antibody (R960-25, Thermo Fisher) used at a concentration of 1:2,000; rabbit α-hexokinase antibody (H2035, US Biological) used at a concentration of 1:5,000-1:10,000; mouse α-GFP JL-8 (cat. no. 632381, Takara) used at a concentration of 1:2,000. Secondary antibodies: goat α-mouse or α-rabbit secondary antibody conjugated to IRDye^®^ 800CW used at a concentration of 1:15,000 (926-32213, LI-COR); α-rabbit secondary conjugated to IRDye^®^ 680RD at a concentration of 1:15,000 (926-68071, LI-COR) Immunoblot quantification was performed by quantifying bands in Image Studio™ (LI-COR). For all blots quantified in this study, Hxk2 loading was normalized to vegetative expression (unless otherwise noted) and raw Sod1-3V5 signal was adjusted based on the corresponding normalized Hxk2 signal.

### Urea denaturation

For the preparation of urea-denatured samples, 3.7-9.25 ODs of cells were pelleted at 1,900 rcf for 2 minutes at 4°C, cells were washed with 1 mL of 10 mM Tris pH 7.5, spun at maximum speed for 1 minute at 4°C, then flash-frozen in liquid nitrogen and stored at -80°C. Thawed pellets were resuspended with 200 µL urea buffer (8M urea in 50mM ammonium bicarbonate) and 200 µL 0.5 mm zirconia/silica beads (cat. no. 11079105z, BioSpec Products). Cells were lysed by bead-beating 4X (5 minutes per cycle) in pre-chilled metal blocks (samples were rested on ice for 2 minutes between cycles), then spun at 20,000 rcf for 10 minutes at 4°C. Supernatant was then transferred to pre-chilled low-adhesion 1.5 mL tubes and samples were spun at 20,000 rcf for 5 minutes at 4°C. After the second spin, supernatants were transferred to pre-chilled low-adhesion 1.5 mL tubes and quantified by Bradford (cat. no. 5000006, Bio-Rad). After quantification, 45 µg of total protein was brought to 60 µL in urea buffer and combined with 30 µL of 3X SDS buffer (18.75 mM Tris pH 6.8, 6% β-mercaptoethanol, 30% glycerol, 9% SDS, 0.05% bromophenol blue). Samples were heated for 5 minutes at 37°C, then stored at -20°C. Urea-denatured samples were run using the same electrophoresis and immunoblot protocol as described for SDS-PAGE.

### Blue native PAGE immunoblotting

Blue native PAGE (BN-PAGE) was performed as outlined by the NativePAGE™ Novex^®^ Bis-Tris Gel System manual (MAN0000557, Life Technologies). 3.7-9.25 ODs of cells were pelleted at 1,900 rcf for 2 minutes at 4°C, cells were washed with 1 mL of 10 mM Tris pH 7.5, spun at maximum speed for 1 minute at 4°C, then flash-frozen in liquid nitrogen and stored at -80°C. To prepare native lysates for BN-PAGE, cell pellets were thawed from -80°C on ice. Thawed pellets were resuspended with 200-300 µL native lysate buffer (1% NP-40 with 1X cOmplete protease inhibitor cocktail [SKU, 11836145001, Roche], 0.5 mM PMSF, and 1% digitonin [SKU D141, Millipore Sigma]) and 200 µL 0.5 mm zirconia/silica beads (cat. no. 11079105z, BioSpec Products). Cell lysis was performed by mechanical disruption using either a FastPrep (MP Biomedicals) or standard bead-beater fitted with pre-chilled metal blocks. Following FastPrep lysis, tubes were punctured using 20.5G needles, then nested in 15 mL conical tubes containing p1000 tips. Samples were spun at 3,000 rpm for 20 seconds at 4°C, then the p1000 tip was used to transfer the flow-through to pre-chilled 1.5 mL low-adhesion tubes. Samples were then spun at 20,000 rcf for 10 minutes at 4°C, then supernatant was transferred to pre-chilled 1.5 mL low-adhesion tubes. An additional spin was performed at 20,000 rcf for 5 minutes at 4°C, then supernatant was transferred to pre-chilled low-adhesion tubes and quantified by Bradford (cat. no. 5000006, Bio-Rad).

Prior to loading, 10-20 µg of total protein of each sample was brought to an equal volume (in native lysis buffer) and 1-2.5 µL of G-250 additive was added to each sample. NativePAGE™ 4-16% Bis-Tris gels (cat. no. BN1004BOX, Invitrogen) were prepared by washing wells 3X with dH_2_O, then 3X with 1X dark blue cathode buffer. After sample loading, gels were run at 150V for 90-120 minutes at 4°C using the anode and cathode buffers outlined in the NativePAGE manual. Transfer to methanol-activated 0.2 µm Immun-Blot^®^ PVDF membranes (cat. no. 1620177, Bio-Rad) was performed at 30V for 1 hour at 4°C in 1X NuPAGE™ buffer (REF NP0006-1, Life Technologies). After transfer, PVDF membranes were incubated in 8% actetic acid for 15 minutes at room temperature. A 5-minute methanol wash was performed to remove excess Coomassie, then membranes were washed 3X in dH_2_O. After this point, blocking, 1° and 2° incubations, and washes were performed as described above for SDS-PAGE immunoblotting.

### Sod in-gel activity assay

Native lysates were prepared following a protocol from Valeria Culotta (Johns Hopkins, Baltimore, MD). 3.7-9.25 ODs of cells were pelleted at 1,900 rcf for 2 minutes at 4°C, cells were washed with 1 mL of 10 mM Tris pH 7.5, spun at maximum speed for 1 minute at 4°C, then flash-frozen in liquid nitrogen and stored at -80°C. To prepare native lysates for activity assays, cells were thawed on ice and resuspended in 200-300 µL native lysis buffer (10 mM NaPi pH 7.8, 0.1% Triton X-100, 5 mM EDTA, 5 mM EGTA, 50 mM NaCl, and 10% glycerol with 1X cOmplete protease inhibitor cocktail [SKU, 11836145001, Roche] and 1 mM PMSF) and 200 µL 0.5 mm zirconia/silica beads (cat. no. 11079105z, BioSpec Products). Cells were lysed by bead-beating 4X (5 minutes per cycle) in pre-chilled metal blocks (samples were rested on ice for 2 minutes between cycles), then spun at 20,000 rcf for 10 minutes at 4°C. Supernatant was then transferred to pre-chilled low-adhesion 1.5 mL tubes and samples were spun at 20,000 rcf for 5 minutes at 4°C. After the second spin, supernatants were transferred to pre-chilled low-adhesion 1.5 mL tubes and quantified by Bradford (cat. no. 5000006, Bio-Rad). Prior to loading, 10-20 µg of total protein of each sample was brought to an equal volume (in native lysis buffer), combined with 5X sample buffer (50% glycerol, 310 mM Tris pH 6.8, 0.5% bromophenol blue; diluted to 1X), then loaded into 4-12% Novex Tris-glycine gels (Invitrogen; wells were washed thoroughly with dH_2_O prior to loading to avoid inhibition of Sod1 activity by sodium azide). Gels were run at 50 mA for 80-90 minutes at 4°C in 1X native gel running buffer (25 mM Tris base, 10 mM EDTA, 192 mM glycine). After running, gels were incubated in 35 mL of Sod staining solution (175 µM nitroblue tetrazolium and 280 µM riboflavin in 0.05M KPi pH 7.8), 35 µL of TEMED (cat. no. 161-0800, Bio-Rad) was added to the staining solution, and gels were incubated in the dark at 4°C for 1 hour. Sod staining solution was then removed, gels were washed 3X in dH_2_O, then incubated in dH_2_O at room temperature in the light to expose bands of Sod enzyme activity. Once sufficiently exposed, gels were placed in plastic sleeve protectors and scanned on a standard color scanner.

### Immunofluorescence staining

To harvest meiotic cells for immunofluorescence, 450 µL aliquots of SPO cultures were added to tubes containing 50 µL of 37% formaldehyde (3.7% final; F79-500, Fisher) and incubated at 4°C overnight. The following day, samples were spun at 1,900 rcf for 2 minutes at room temperature, washed with 1 mL 0.1M KPi pH 6.4, resuspended in 1 mL 1.2M sorbitol-citrate (100 µM K_2_HPO_4_, 36 µM citric acid, 1.2M D-sorbitol), and stored at -20°C. To permeabilize cells, samples were thawed at room temperature, then spheroplasted in digestion mix (8.9% glusulase [part no. NEE154001EA, PerkinElmer] and 1 mg/mL 100T zymolyase [cat. no. 08320932, MP Biomedicals or cat. no. 320932, VWR] in 1.2M sorbitol-citrate) by rotating at 30°C for 4-5 hours (digestion was assessed by looking at cells on a light microscope). After spheroplasting, cells were spun at 900 rcf for 2 minutes, resuspended in 1 mL 1.2M sorbitol-citrate, spun at 1,900 rcf for 2 minutes, then resuspended in a variable volume (typically 25-100 µL) of 1.2M sorbitol-citrate to achieve satisfactory cell density for immunofluorescence. Prior to the addition of cells, wells of frosted microscope slides (design 161-041-122, TEKDON, Inc.) were treated with 0.1% (w/v) poly-L-lysine solution (cat. no. P8920, Sigma) for 5 minutes at room temperature, excess poly-L-lysine was aspirated off, and wells were allowed to air dry. To prepare cells for immunofluorescence, 5 µL of digested cell solutions were pipetted onto poly-L-lysine-treated wells and incubated at room temperature for 10 minutes. Excess cell solution was then aspirated off and cell density was checked under a light microscope. Slides were incubated in cold methanol for 3 minutes, then placed directly in cold acetone for 10 seconds. After acetone incubation, slides were allowed to dry completely prior to addition of primary antibody solution.

For Sod1-3V5 immunofluorescence staining, 1:6 pre-absorbed mouse α-V5 primary antibody (R960-25, Thermo Fisher) was diluted to 1:800 in 1X PBS-BSA, 5 µL of primary dilution was added to each well, and slides were incubated at 4°C overnight. The next day, wells were washed 3X with 1X PBS-BSA, 5 µL of secondary antibody solution (1:6 pre-absorbed α-mouse Cy3, diluted to 1:200 in 1X PBS-BSA; 115-165-003, Jackson ImmunoResearch Laboratories) was added to each well, and samples were incubated in the dark for 2.5 hours at room temperature.

For tubulin immunofluorescence staining, rat α-tubulin primary antibody (RID AB_325005, MCA78G, Bio-Rad) was diluted 1:200 in 1X PBS-BSA, 5 µL of primary dilution was added to each well, and slides were incubated at 4°C overnight. The next day, wells were washed 2X with 1X-PBS-BSA, 5 µL of secondary antibody solution (1:6 pre-absorbed α-rat FITC, diluted to 1:200 in PBS-BSA; 712-095-153, Jackson ImmunoResearch Laboratories) was added to each well, and samples were incubated in the dark for one hour at room temperature.

After 2-3X with 1X PBS-BSA (3X for Sod1-3V5 staining and 2X for tubulin staining), a drop of VECTASHIELD^®^ with DAPI (REF H-1200, Vector Laboratories, Inc.) was added to each well before coverslip addition. Slides were sealed with clear nail polish and dried for at least 10 minutes prior to imaging or storage at -20°C. For Sod1 IF analysis, all information (strain, time point, DAPI staining, genotype) was blinded during scoring of Sod1-3V5 IF foci. In Fiji, cells were numbered and scored as containing dim, moderate, or bright foci. Staging of cells was assessed by DAPI staining.

### Fluorescence microscopy

All fluorescence microscopy in this study was performed using a DeltaVision Elite microscope (GE Healthcare) using either a 60X/1.42 NA oil-immersion objective (for tubulin and DAPI staining) or 100X/1.40 NA oil-immersion objective (for Sod1 immunofluorescence) objective. After acquisition, images were deconvolved using softWoRx imaging software (GE Healthcare).

## Data availability

All reagents (strains or plasmids) used in this study are available upon request.

## Acknolwedgements

We thank members of the Brar and Ünal labs for their feedback on the manuscript, as well as for help with experimental design and technical support. We also thank Valeria Culotta for sharing protocols and reagents, and Amit Reddi for manuscript feedback.

## SUPPLEMENTAL FIGURE LEGENDS

**Figure S1.** (A) Mitotic growth of wild-type and *sod1*Δ diploids. To assess mitotic growth, wild-type and *sod1*Δ strains were serial-diluted and plated on rich media (1X YEP with 2% dextrose). From left to right, OD_600_ values plated were 0.2, 0.04, 0.008, 0.0016, and 0.00032. Image shows growth after 72 hours at 30°C.

**Figure S2.** (A) *SOD1* mRNA expression under various stress conditions. Northern blotting for *SOD1^LUTI^* and *SOD1^canon^*. under stress conditions during vegetative exponential growth (MB = methylene blue). From left to right: untreated (untr.), 5 mM DTT (DTT), 1.5 mM diamide (DA), 1 mM paraquat (PQ), 0.3 mM hydrogen peroxide (H_2_O_2_), and 37°C heat shock (37°C). Conditions tested were based on Gasch et al., 2000, and samples were harvested 1 hour after treatment. (B) *SOD1^LUTI^* and *HNT1^LUTI^* expression in wild-type and UPRE mutant cells upon UPR^ER^ activation. RT-qPCR analysis of *SOD1^LUTI^* and *HNT1^LUTI^* levels 15-120 minutes after treatment with 5 mM DTT. For RT-qPCR, cDNA was prepared from total RNA samples used in Figure 3D, and fold-change (2^−ΔC_T_^) values were calculated using *ACT1* mRNA as a control. *HNT1^LUTI^* is a positive control for UPR^ER^ induction (Van Dalfsen et al., 2018).

**Figure S3.** (A) Total Sod1 protein decreases in meiotic, urea-denatured lysates. SDS-PAGE and immunoblotting for Sod1-3V5 in lysates prepared with 8M urea (quantification shown below). (B) Meiotic Sod1 levels in wild-type and LUTI-disrupted strains. SDS-PAGE immunoblot quantification of Sod1 protein levels in synchronized cells with or without (+ *CYC1t*) LUTI expression (\**pGAL*-*NDT80* release at 5 hours). Average values were calculated for each genotype in triplicate (paired t test two-tailed P = 0.0005).

**Figure S4.** (A) Blue native PAGE (BN-PAGE) of Sod1-3V5 in meiosis. Soluble, dimeric Sod1 (32 kDa) detected in vegetative and meiotic native lysates (\**pGAL*-*NDT80* release at 5 hours). (B) Vacuolar protease activity is not responsible for the loss of dimeric Sod1 from native meiotic lysates. Vegetative native lysate was incubated in either native buffer or meiotic lysate (prepared from meiotic sample harvested after 7 hours in sporulation media) and either preincubated on ice for 30 minutes (+) or prepared fresh prior to loading (–). (C) Soluble, dimeric Sod1 is generated from a *pATG8*-*SOD1*-*3V5* transgene. BN-PAGE (top) and SDS-PAGE (bottom) probing for Sod1-3V5 produced from a *pATG8*-driven transgene. (D) Sod enzymatic activity during meiosis. In-gel activity assay probing for Sod1 (top band) and Sod2 (bottom bands) activity in native lysates (\**pGAL*-*NDT80* release at 5 hours).

**Figure S5.** (A) ALS-associated mutant Sod1 decreases sporulation efficiency and spore viability. To assess spore viability, 80 cells (20 tetrads) were dissected per genotype on YEP +2% dextrose and incubated at 30°C for 48 hours. (B) Expression of *pATG8-SOD1* transcripts. RT-qPCR analysis of wild-type, G92A, and A3V *pATG8*-*SOD1* transcripts. For RT-qPCR, cDNA was prepared from total RNA samples matching SDS-PAGE samples shown in Figure 7B, and fold-change (2^−ΔC_T_^) values were calculated using *PFY1* mRNA as a control. To detect *pATG8*-*SOD1* mRNA specifically, expression was assessed in *sod1*Δ strains. (C) *pATG8-hSOD1* transcript and protein expression. Matched RT-qPCR (left) and SDS-PAGE immunoblot (right) showing expression from a *pATG8*-*hSOD1* (yeast codon-optimized) transgene. For RT-qPCR, fold-change (2^−ΔC_T_^) values were calculated using *PFY1* mRNA as a control. Expression was assessed in a *sod1*Δ strain. (D) Sod1^WT^-GFP and Sod1^G92A^-GFP localization in fixed meiotic cells. Wild-type and G92A localization in cells fixed after 0, 2, and 4 hours in sporulation media. Identical exposure conditions were used during image acquisition, but post-acquisition exposures are different for wild-type and G92A micrographs to improve the visibility of G92A protein. (scale bars = 5 µm). (E) Wild-type and G92A spore viability with 3V5 and GFP tags. For each genotype, 80 cells (20 tetrads) were dissected on YEP +2% dextrose and incubated at 30°C for 48 hours.

## Notes

### Competing Interest Statement

The authors have declared no competing interest.

