## Supplementary figures and images for "Meiotic resetting of the cellular Sod1 pool is driven by protein aggregation, degradation, and transient LUTI-mediated repression"

### Supplemental Figures

# A

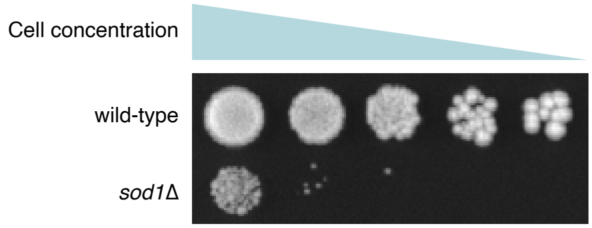


**Figure S1.**

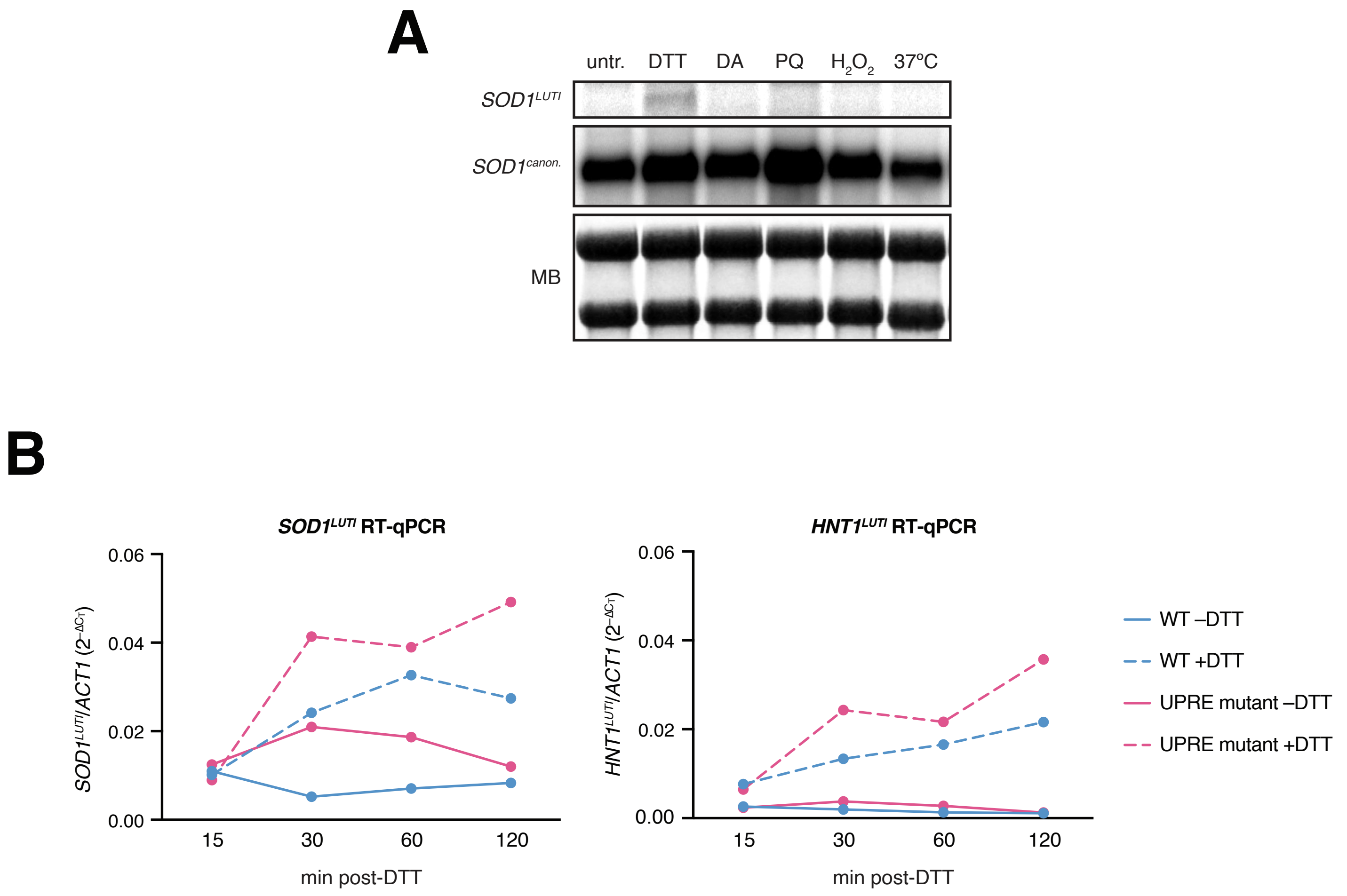

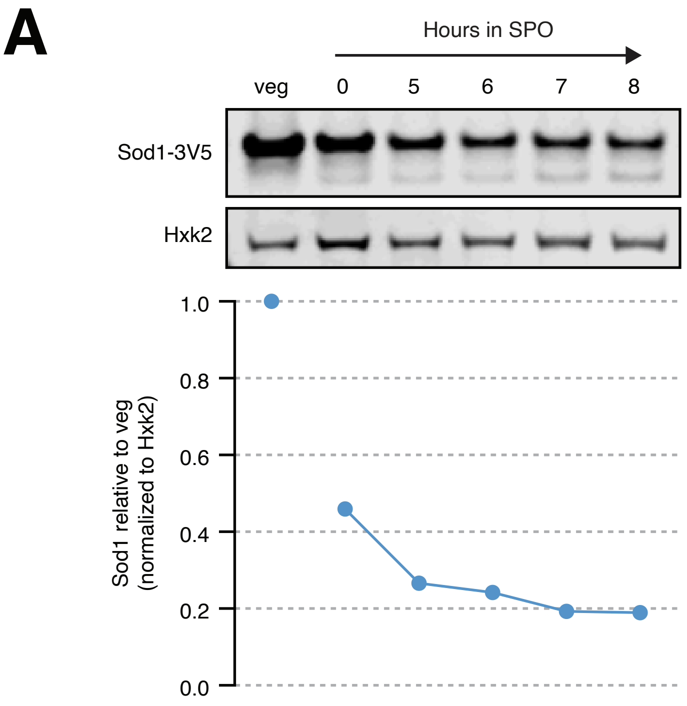

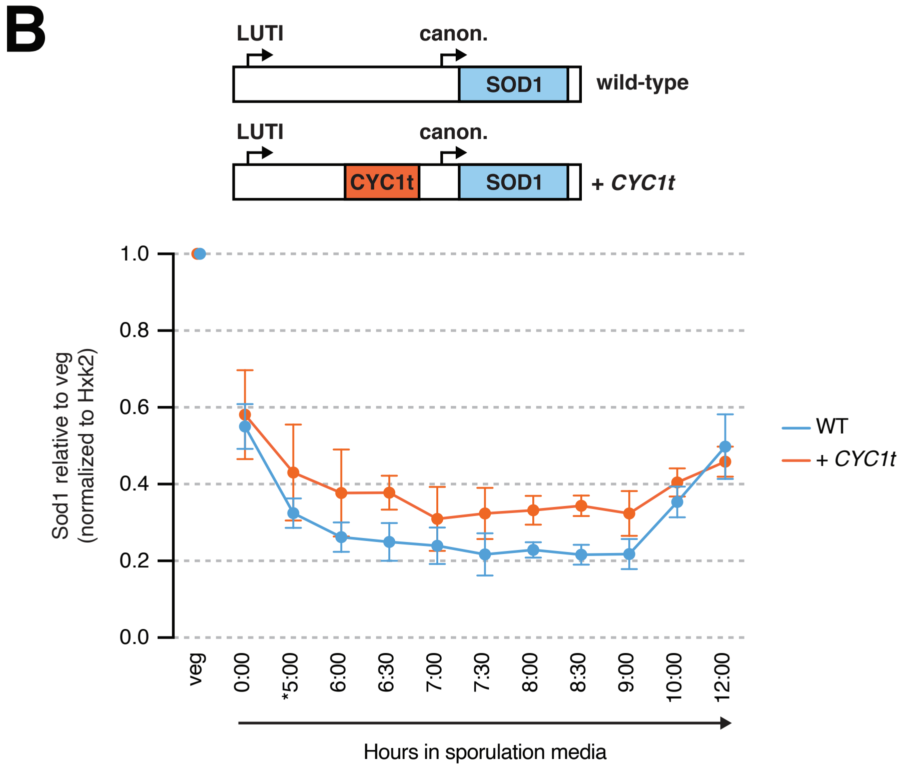

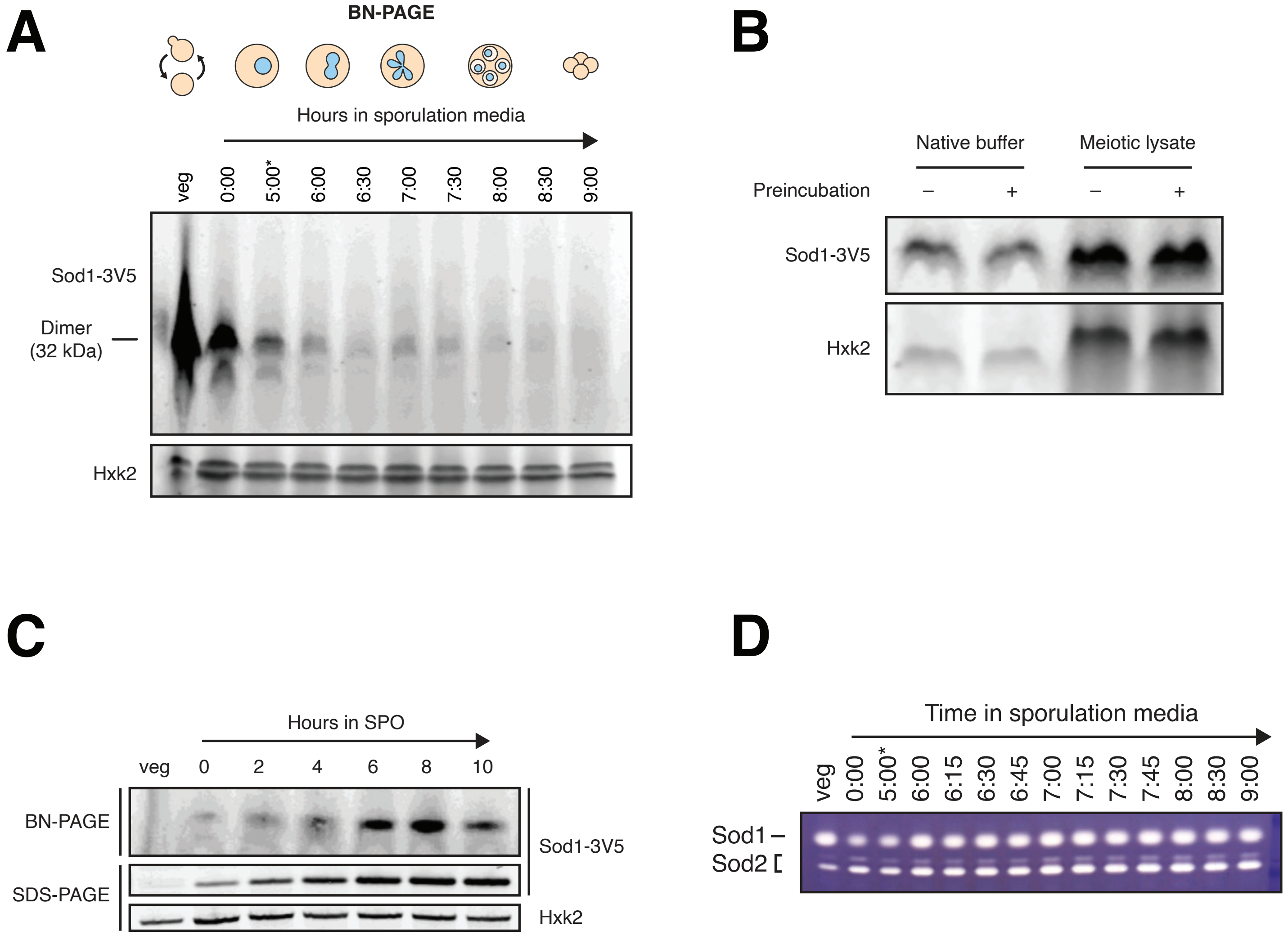

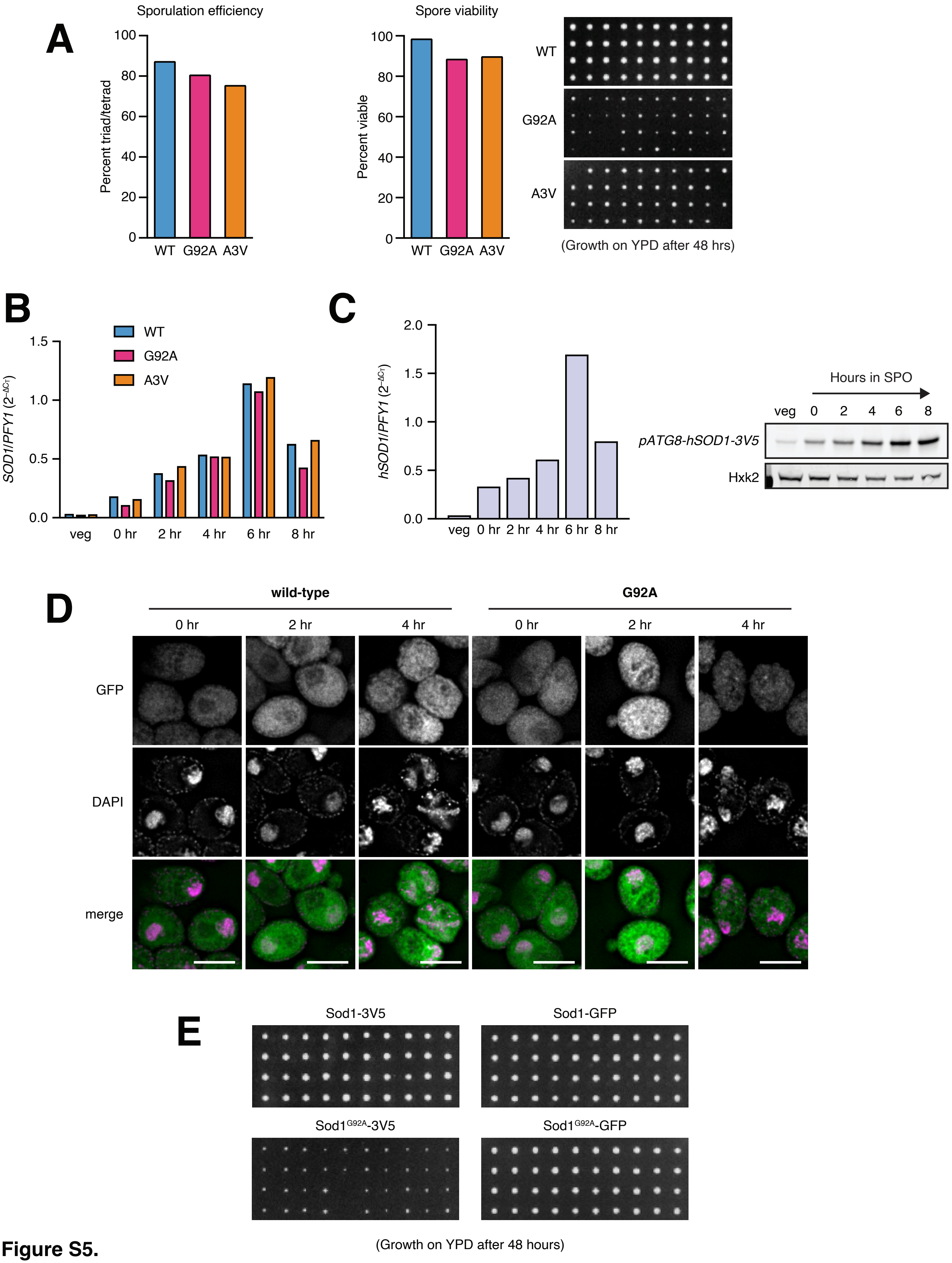
