## Supplemental Table 1 for "Meiotic resetting of the cellular Sod1 pool is driven by protein aggregation, degradation, and transient LUTI-mediated repression"

**Table S1. Strain table.**

| **Strain** | **Genotype** |
| --- | --- |
| SK1 wild-type | *ho::LYS2 lys2 ura3 leu2::hisG his3::hisG trp1::hisG* |
| ÜB13 | MAT**a** |
| ÜB15 | MAT**a**/MATalpha |
| ÜB95 | MAT**a**/MATalpha  GAL-NDT80::TRP1/GAL-NDT80::TRP1  ura3::pGPD1-GAL4(848).ER::URA3/ura3::pGPD1-GAL4(848).ER |
| ÜB1362 | MAT**a**/MATalpha  TRP1/TRP1  HIS3/HIS3  HIS4/HIS4  REC8-3HA::URA3/REC8-3HA::URA3  lys2::TetOx240:URA3/lys2::TetOx240:URA3  leu2::LEU2 tetR-GFP/leu2::LEU2 tetR-GFP |
| ÜB4431 | MAT**a**/MATalpha  TRP1/TRP1  HIS3/HIS3  HIS4/HIS4  REC8-3HA::URA3/REC8-3HA::URA3  lys2::TetOx240:URA3/lys2::TetOx240:URA3  leu2::LEU2 tetR-GFP/leu2::LEU2 tetR-GFP  hac1::NatMX6/hac1::KanMX6 |
| ÜB9084 | MAT**a**/MATalpha  sod1::KanMX6/sod1::KanMX6 |
| ÜB15290 | MAT**a**/MATalpha  GAL-NDT80::TRP1/GAL-NDT80::TRP1  ura3::pGPD1-GAL4(848).ER::URA3/ura3::pGPD1-GAL4(848).ER  Sod1-3V5::KanMX6/Sod1-3V5::KanMX6 |
| ÜB24294 | MAT**a**/MATalpha  GAL-NDT80::TRP1/GAL-NDT80::TRP1  ura3::pGPD1-GAL4(848).ER::URA3/ura3::pGPD1-GAL4(848).ER  Sod1-3V5::KanMX6/Sod1-3V5::KanMX6  CYC1t (unmarked)/CYC1t (unmarked) |
| ÜB25370 | MAT**a**/MATalpha  Sod1-3V5::KanMX6/Sod1-3V5::KanMX6 |
| ÜB26718 | MAT**a**/MATalpha  his3::HIS3::pATG8-SOD1-3V5/his3::HIS3::pATG8-SOD1-3V5 |
| ÜB27568 | MAT**a**/MATalpha  pdr5::hygBMX6/pdr5::hygBMX6  his3::HIS3::pATG8-SOD1-3V5/his3::HIS3::pATG8-SOD1-3V5 |
| ÜB27570 | MAT**a**/MATalpha  pdr5::hygBMX6/pdr5::hygBMX6  his3::HIS3::pATG8-SOD1^G92A^-3V5/his3::HIS3::pATG8-SOD1^G92A^-3V5 |
| ÜB28898 | MAT**a**/MATalpha  Sod1^G92A^-3V5::KanMX6/ Sod1^G92A^-3V5::KanMX6 |
| ÜB29380 | MAT**a**/MATalpha  sod1::KanMX6/sod1::KanMX6  his3::HIS3::pATG8-SOD1-3V5/his3::HIS3::pATG8-SOD1-3V5 |
| ÜB29382 | MAT**a**/MATalpha  sod1::KanMX6/sod1::KanMX6  his3::HIS3::pATG8-SOD1^G92A^-3V5/his3::HIS3::pATG8-SOD1^G92A^-3V5 |
| ÜB29584 | MAT**a**/MATalpha  Sod1-yoEGFP::KanMX6/Sod1-yoEGFP::KanMX6 |
| ÜB29894 | MAT**a**/MATalpha  Sod1^G92A^-yoEGFP::KanMX6/Sod1^G92A^-yoEGFP::KanMX6 |
| ÜB30400 | MAT**a**/MATalpha  sod1::KanMX6/sod1::KanMX6  his3::HIS3::pATG8-SOD1^A3V^-3V5/his3::HIS3::pATG8-SOD1^A3V^-3V5 |
| ÜB30402 | MAT**a**/MATalpha  sod1::KanMX6/sod1::KanMX6  his3::HIS3::pATG8-hSOD1-3V5/his3::HIS3::pATG8-hSOD1-3V5 |
| ÜB31016 | MAT**a**/MATalpha  pdr5::hygBMX6/pdr5::hygBMX6  CYC1t (unmarked)/CYC1t (unmarked)  Sod1-3V5::KanMX6/Sod1-3V5::KanMX6 |
| ÜB31129 | MAT**a**  SOD1-LUTI-distal-UPRE-scramble (unmarked)  SOD1-LUTI-proximal-UPRE∆ (unmarked) |
| ÜB31188 | MAT**a**/MATalpha  pdr5::hygBMX6/pdr5::hygBMX6  Sod1-3V5::KanMX6/Sod1-3V5::KanMX6 |
| ÜB32502 | MAT**a**/MATalpha  Sod1^A3V^-3V5::KanMX6/Sod1^A3V^-3V5::KanMX6 |
| ÜB32660 | MAT**a**/MATalpha  Ime1::HygB/ime1::HygB  Sod1-3V5::KanMX6/Sod1-3V5::KanMX6 |
| ÜB32700 | MAT**a**/MATalpha  trp1::pGPD1-LexA-ER-HA-B112::TRP1/trp1::pGPD1-LexA-ER-HA-B112::TRP1  KanMX:p8X-LexO-pCyc1-SOD1-LUTI/KanMX:p8X-LexO-pCyc1-SOD1-LUTI  Sod1-3V5 (unmarked)/ Sod1-3V5 (unmarked) |
| ÜB32702 | MAT**a**/MATalpha  trp1::pGPD1-LexA-ER-HA-B112::TRP1/trp1::pGPD1-LexA-ER-HA-B112::TRP1  KanMX:p8X-LexO-pCyc1-SOD1-LUTI/KanMX:p8X-LexO-pCyc1-SOD1-LUTI  CYC1t (unmarked)/CYC1t (unmarked)  Sod1-3V5 (unmarked)/ Sod1-3V5 (unmarked) |
