## Supplemental Table 2 for "Meiotic resetting of the cellular Sod1 pool is driven by protein aggregation, degradation, and transient LUTI-mediated repression"

**Table S2. Primers used for strain construction.**

Lowercase letters = annealing region to template

**Pringle strain construction**

| **Construct name** | **Forward primer (5′ to 3′)** | **Reverse primer (5′ to 3′)** |
| --- | --- | --- |
| *SOD1*-*3V5* | CGGTCCAAGACCAGCCTGTGGTGTCATTGGTCTAACCAACcggatccccgggttaattaa | ACTTACATACGGTTTTTATTCAAGTATATTATCATTAACAgaattcgagctcgtttaaac |
| *SOD1*-*yoEGFP* | CGGTCCAAGACCAGCCTGTGGTGTCATTGGTCTAACCAACggtgacggtgctggttta | ACTTACATACGGTTTTTATTCAAGTATATTATCATTAACAgaattcgagctcgtttaaac |
| *sod1*∆ | AAAGCAATCGCGCAGACAAATAAAACATAATTAATTTATAcggatccccgggttaattaa | ACTTACATACGGTTTTTATTCAAGTATATTATCATTAACAgaattcgagctcgtttaaac |
| *8XlexO*-*LUTI* | AGCGCCTCTTTTCCTTATTGTGGTAACGGTGGCTTATTTGgaattcgagctcgtttaaac | CTTTTCATTCGTAGACATGTTTACCGCTTGTTCTTAAACaagcttgatatcgaattcctg |
| *ime1*∆ | ATAAAAGAAAAGCTTTTCTATTCCTCTCCCCACAAACAAAcggatccccgggttaattaa | TTGAGGGAAGGGGGAAGATTGTAGTACTTTTCGAGAAcactagtggatctgatatcatcg |

**Cas9 strain construction**

Blue = PAM mutations

Red = coding changes

Purple = PAM/coding mutation

| **Construct name** | **Repair top (5′ to 3′)** | **Repair bottom (5′ to 3′)** |
| --- | --- | --- |
| *SOD1-3V5* | CGGTCCAAGACCAGCCTGTGGTGTCATTGGTCTAACTAACcggatccccgggttaattaa | ACTTACATACGGTTTTTATTCAAGTATATTATCATTAACAttatttagaagtggcgcgcc |
| *CYC1t* | TATCTCTGAAGTGCAGCCGATTGGGCGTGCGACTCACCCAtcatgtaattagttatgtca | CTGTCATCGCCTTATCGAACCCGCTACTGAGATCATGTCGTGAGgcaaattaaagccttc |
| Sod1^G92A^ | TCCAACTGACGAAGTCAGACATGTCGGTGACATGGGTAACgtaaagacggacgaaaatgc | AAGCTTGATCAAAGAGTCCTTGAAGGAGCCCTTGGCCACAGcattttcgtccgtctttac |
| Sod1^A3V^ | CAAATAAAACATAATTAATTTATAATGGTTCAAGTAGTCGcagtgttaaagggtgatgca | ATTCGGAAGCCTGTTCGAACTTGACAACACCAGAGACACCtgcatcaccctttaacactg |
| UPRE^prox^∆ | TGGTTTAAGAACAAGCGGTAAACATGTCTACGAATGAAAAGCTAGAGCAAagagtatatt | TAAATAGATTCTTCTTACAATATAATAGAAATAAAATATACTCTttgctctagcttttca |
| UPRE^dist^ scramble | AAGCTCGGCACAATAAACGTTACTGAAAgtttaggatctttgtaaagtaacctaggatgg | ATAATATGGACTGGGCAATCAATGTCGCccatcctaggttactttacaaagatcctaaac |
