## Supplemental Table 3 for "Meiotic resetting of the cellular Sod1 pool is driven by protein aggregation, degradation, and transient LUTI-mediated repression"

**Table S3. Primers for northern blotting, RT-qPCR, and smRNA-FISH**

**Northern blot primers**

| **Primer** | **Sequence (5**′ **to 3**′**)** |
| --- | --- |
| *SOD1* F | CAACCACTGTCTCTTACGAGATCGC |
| *SOD1* R | taatacgactcactataggCACCATTTTCGTCCGTCTTTACG  (lowercase letters denote T7 promoter) |

**RT-qPCR primers**

| **Primer** | **Sequence (5**′ **to 3**′**)** |
| --- | --- |
| *SOD1* F | CTCTTTGATCAAGCTTATCGGTCC |
| *SOD1* R | AGATTCTTCAGTGTCACCCTTACC |
| *hSOD1* F | GACTCCGTAATCAGCCTTTCAG |
| *hSOD1* R | TCTCCTCGTTTCCTCCCTTC |
| *SOD1^LUTI^* F | TATTGGCATGATGCGAAATTGGAC |
| *SOD1^LUTI^* R | ATCGAACCCGCTACTGAGATC |
| *HNT1^LUTI^* F | TGGTGCGAATCGTTACAGAA |
| *HNT1^LUTI^* R | AATGCTTCAGTAGGGCGGTA |
| *ACT1* F | GTACCACCATGTTCCCAGGTATT |
| *ACT1* R | AGATGGACCACTTTCGTCGT |
| *PFY1* F | acggtagacatgatgctgagg |
| *PFY1* R | acggttggtggataatgagc |

**smRNA-FISH probes**

| **Probe** | **Target** | **Sequence (5′ to 3′)** |
| --- | --- | --- |
| 1 | *SOD1* ORF | ttaacactgcgactgcttga |
| 2 | *SOD1* ORF | tgacaacaccagagacaccg |
| 3 | *SOD1* ORF | gattcggaagcctgttcgaa |
| 4 | *SOD1* ORF | cgtaagagacagtggttggc |
| 5 | *SOD1* ORF | ttaggactgttaccagcgat |
| 6 | *SOD1* ORF | gaatatggaacccacgttct |
| 7 | *SOD1* ORF | cattggtggcatctccaaac |
| 8 | *SOD1* ORF | aagtgaggaccagcagagac |
| 9 | *SOD1* ORF | gtgtgtcttcttgaaaggat |
| 10 | *SOD1* ORF | tgacttcgtcagttggagca |
| 11 | *SOD1* ORF | ttacccatgtcaccgacatg |
| 12 | *SOD1* ORF | accattttcgtccgtcttta |
| 13 | *SOD1* ORF | aaagagtccttgaaggagcc |
| 14 | *SOD1* ORF | acggaggtaggaccgataag |
| 15 | *SOD1* ORF | cgtggataacgacgcttctg |
| 16 | *SOD1* ORF | cttacctaagtcatcttggc |
| 17 | *SOD1* ORF | ggcattaccagtcttcaaag |
| 18 | *SOD1* ORF | tagaccaatgacaccacagg |

| **Probe** | **Target** | **Sequence** |
| --- | --- | --- |
| 1 | 5' LUTI extension | ttttatggctgggttgacta |
| 2 | 5' LUTI extension | cagtcaaacctttggactct |
| 3 | 5' LUTI extension | tgtcgaaccaacagtggtcg |
| 4 | 5' LUTI extension | gaattgtcaaggactctcca |
| 5 | 5' LUTI extension | gatttggaagctattgctca |
| 6 | 5' LUTI extension | ttccaattagactgtgccaa |
| 7 | 5' LUTI extension | ctcatgtcttcaaagacgca |
| 8 | 5' LUTI extension | aactttagtgctcctttcat |
| 9 | 5' LUTI extension | tatcgttaagttgggccaac |
| 10 | 5' LUTI extension | tcttctacatcggtggtttc |
| 11 | 5' LUTI extension | gacttggtcgaagcagctaa |
| 12 | 5' LUTI extension | agaccacttggacaagcatt |
| 13 | 5' LUTI extension | tattggtaccggtaagtgtg |
| 14 | 5' LUTI extension | tggtgtcaagtctatgtacc |
| 15 | 5' LUTI extension | cagcgagagattgcttaacg |
| 16 | 5' LUTI extension | gttccgtcggtaaggacaag |
| 17 | 5' LUTI extension | aaaacactgctagaggggca |
| 18 | 5' LUTI extension | acaaagcttgttgcaggtgg |
| 19 | 5' LUTI extension | atcaggcgatgctaagatgg |
| 20 | 5' LUTI extension | acgacgcaattttggtcgat |
| 21 | 5' LUTI extension | tttagatttccaagccgacg |
| 22 | 5' LUTI extension | gagcaagcaattatgaccgc |
| 23 | 5' LUTI extension | ggtagtcatctcagtttatc |
| 24 | 5' LUTI extension | tataggcatgccatgtctaa |
| 25 | 5' LUTI extension | attattcttcgagcccaatt |
| 26 | 5' LUTI extension | tgtcgtcgtacgtgatgagg |
| 27 | 5' LUTI extension | ttttcacatggatgagtcgc |
| 28 | 5' LUTI extension | aactttttttgcagatgccg |
| 29 | 5' LUTI extension | gccaatattacctgtggaag |
| 30 | 5' LUTI extension | cttacgtccaatttcgcatc |
| 31 | 5' LUTI extension | aatcggctgcacttcagaga |
| 32 | 5' LUTI extension | ctactgagatcatgtcctga |
| 33 | 5' LUTI extension | ctgtcatcgccttatcgaac |
| 34 | 5' LUTI extension | ggttctgtacttccagtaag |
| 35 | 5' LUTI extension | cgcttactaattccgacaca |
| 36 | 5' LUTI extension | tatatatctgcgatggcgag |
