## Supplemental Table 4 for "Meiotic resetting of the cellular Sod1 pool is driven by protein aggregation, degradation, and transient LUTI-mediated repression"

**Table S4. Plasmids used for strain construction.**

| **Plasmid Name** | **Description** |
| --- | --- |
| pÜB1 | pFA6a-KanMX6 |
| pÜB81 | pFA6a-3V5-KanMX6 |
| pÜB182 | pFA6a-yoEGFP-KanMX6 |
| pÜB196 | p415-GalL-Cas9-CYC1t |
| pÜB217 | pFA6a-hphNT1-HygR |
| pÜB925 | pL10-p8LexOCYC1 |
| pÜB1347 | CEN/ARS-URA3-pPGK1-Cas9-SOD1-LUTI-proximal-UPRE-gRNA  (gRNA sequence = 5’-GTCTACGAATGAAAAGCTAG-3’) |
| pÜB1372 | CEN/ARS-URA3-pPGK1-Cas9-SOD1-G92-gRNA  (gRNA sequence = 5’-AACGTAAAGACGGACGAAAA-3’) |
| pÜB1641 | CEN/ARS-URA3-pPGK1-Cas9-SOD1-LUTI-disruption-gRNA  (gRNA sequence = 5’-GGCGTGCGACTCACCCACTC-3’) |
| pÜB1941 | HIS3-pATG8-SOD1-3V5 |
| pÜB1942 | HIS3-pATG8-SOD1^G92A^-3V5 |
| pÜB2069 | CEN/ARS-URA3-pPGK1-Cas9-SOD1-LUTI-distal-UPRE-gRNA  (gRNA sequence = 5’-CCATCCACGATTACTTTACA-3’) |
| pÜB2070 | HIS3-pATG8-SOD1^A3V^-3V5 |
| pÜB2116 | HIS3-pATG8-hSOD1-3V5 |
| pÜB2237 | CEN/ARS-URA3-pPGK1-Cas9-SOD1-A3-gRNA  (gRNA sequence = 5’-CTTGACAACACCAGAGACAC-3’) |
| pÜB2257 | CEN/ARS-URA3-pPGK1-Cas9-SOD1-C-term-gRNA  (gRNA sequence = 5’-ATATTATCATTAACATTAGT-3’) |
